# Active site remodeling in tumor-relevant IDH1 mutants drives distinct kinetic features and potential resistance mechanisms

**DOI:** 10.1101/2024.01.10.574970

**Authors:** Matthew Mealka, Nicole A. Sierra, Diego Avellaneda Matteo, Elene Albekioni, Rachel Khoury, Timothy Mai, Brittany M. Conley, Nalani J. Coleman, Kaitlyn A. Sabo, Elizabeth A. Komives, Andrey A. Bobkov, Andrew L. Cooksy, Steve Silletti, Jamie M. Schiffer, Tom Huxford, Christal D. Sohl

## Abstract

Mutations in human isocitrate dehydrogenase 1 (IDH1) drive tumor formation in a variety of cancers by replacing its conventional activity with a neomorphic activity that generates an oncometabolite. Little is understood of the mechanistic differences among tumor-driving IDH1 mutants. We previously reported that the R132Q mutant uniquely preserves conventional activity while catalyzing robust oncometabolite production, allowing an opportunity to compare these reaction mechanisms within a single active site. Here, we employed static and dynamic structural methods and found that, compared to R132H, the R132Q active site adopted a conformation primed for catalysis with optimized substrate binding and hydride transfer to drive improved conventional and neomorphic activity over R132H. This active site remodeling revealed a possible mechanism of resistance to selective mutant IDH1 therapeutic inhibitors. This work enhances our understanding of fundamental IDH1 mechanisms while pinpointing regions for improving inhibitor selectivity.

## Introduction

Wild type (WT) IDH1 is a highly conserved cytosolic and peroxisomal homodimeric enzyme that reversibly converts isocitrate (ICT) to α-ketoglutarate (αKG) in a NADP^+^-dependent oxidative decarboxylation. However, tumor-driving IDH1 mutants catalyze the NADPH-dependent conversion of αKG to the oncometabolite D-2-hydroxyglutarate (D2HG), while also typically ablating the conventional reaction ^1–3^. D2HG competitively inhibits αKG-dependent enzymes like TET2 and JmjC lysine demethylases to cause DNA and histone hypermethylation and cellular de-differentiation ^4,5^. Mutations at R132 drive >85% lower grade and secondary gliomas ^6^ and ∼40% of cartilaginous tumors ^7^, with R132H typically the most common ^8,9^. These mutated enzymes have been successfully therapeutically targeted, with several FDA-approved allosteric selective inhibitors in use and more in clinical trials (recently reviewed in ^10–12^).

While early kinetic characterization of IDH focused on bacterial forms, recent efforts have illuminated details of human IDH1. As IDH1 WT binds its substrates, a conformational change occurs where the large domain (residues 1-103 and 286-414) and small domain (residues 104-136 and 186-285) move towards each other relative to a hinge point (residues 134-141) in the clasp domain (residues 137-185) ^13^. This movement closes the active site cleft with a concomitant opening of a back cleft ^13^. A critical regulatory segment comprised of the α10 helix (residues 271-285) helps stabilize an open, inactive conformation ^13^ in the absence of substrates, and undergoes a conformational change to help properly orient the active site residues upon substrate binding-driven closure ^13^. These structural features are generally preserved in IDH1 R132H ^1,3,14^, but inherent catalytic deficiencies coupled with improved NADPH binding allow this mutant to catalyze inefficient D2HG production, albeit at great benefit to the tumor environment.

To better understand how D2HG production occurs, there is tremendous value in studying a mutant with more robust neomorphic reaction catalytic efficiency. IDH1 R132S/L/G/Q mutations have also been reported in patients with various frequencies ^15–19^, and different mutations have been shown to support distinct D2HG levels in tumors ^20^. We have shown that these mutants have unique kinetic properties for both neomorphic and conventional reactions ^21,22^, suggesting that their kinetic features may drive some of the variability in patients’ D2HG levels ^22^. We identified one mutant, R132Q, that uniquely maintained WT-like properties with modest conventional catalytic activity, but also drove unusually robust D2HG production ^21^ and was resistant to mutant IDH1 inhibitors via a mechanism not yet understood ^22^. This R132Q mutant has been shown to drive enchondroma tumor formation in mouse models ^23^. By establishing unique features of IDH1 R132Q and R132H mutants, we can identify additional selectivity handles for improved mutant IDH1 inhibitors, as an H-to-Q mutation requires only a single base change. Investigating the atomic-level mechanisms that drive such diverse kinetic activity and inhibition among tumor-relevant IDH1 mutants can also inform the types of chemical features that can guide the field of enzyme design ^24^.

Here, we establish the static and dynamic structural features that drive the unique kinetic properties among tumor-relevant IDH1 mutants, capitalizing on the unusual active site attributes that allow IDH1 R132Q to maintain both normal and neomorphic activities. We identified multiple conformations for R132Q binding to the neomorphic substrate αKG, but not the conventional substrate ICT. Our kinetics and dynamic structural methods clarified that R132Q’s ability to explore multiple conformations and substrate binding modes occurred within a relatively immobile, solvent-inaccessible enzyme that is better optimized for substrate binding, hydride transfer, and mutant IDH1 inhibitor resistance as compared to R132H.

## Results

### Kinetic features of IDH1 R132Q suggest structurally optimized substrate binding and hydride transfer steps relative to R132H

We previously demonstrated that IDH1 R132Q uniquely maintains modest catalytic efficiency for the conventional reaction (ICT to αKG conversion), while also displaying much higher catalytic efficiency for the neomorphic reaction (αKG to D2HG conversion) relative to R132H ^21,22^. Steady-state kinetics analysis performed for this present study (Extended Data Fig. 1) revealed a 5.6-fold increase in catalytic efficiency for the conventional reaction in IDH1 R132Q versus R132H, driven primarily by an increase in *k*_cat_. R132Q catalyzed the neomorphic reaction 9.4-fold more efficiently than R132H via an increase in *k*_cat_ and decrease in *K*_m_. This suggests that IDH1 R132Q exhibits a more stable transition state and provides more optimized on/off paths of the reactants and products compared to R132H.

Pre-steady-state kinetics experiments indicated that hydride transfer, or a step preceding it, was rate-limiting for the conventional reaction catalyzed by WT and R132Q, and for the neomorphic reaction catalyzed by R132Q and R132H (Fig. 1). Interestingly, NADPH consumption by R132H showed an initial lag that could be eliminated upon using higher concentrations of αKG (Extended Data Fig. 2). A lag has been reported previously with IDH1 WT, which was eliminated via pre-incubation of both ICT and metal ^25–28^. Interestingly, we did not observe a lag in the neomorphic reaction catalyzed by R132Q, despite using a concentration of αKG that was 10-fold lower than the concentration associated with a lag in R132H. This suggests that αKG is more proficient at driving IDH1 R132Q from an inactive to an active state compared to R132H, though it was not apparent through these experiments whether this was achieved by a more catalytically primed ground state or a faster conformational change.

**Fig. 1.**
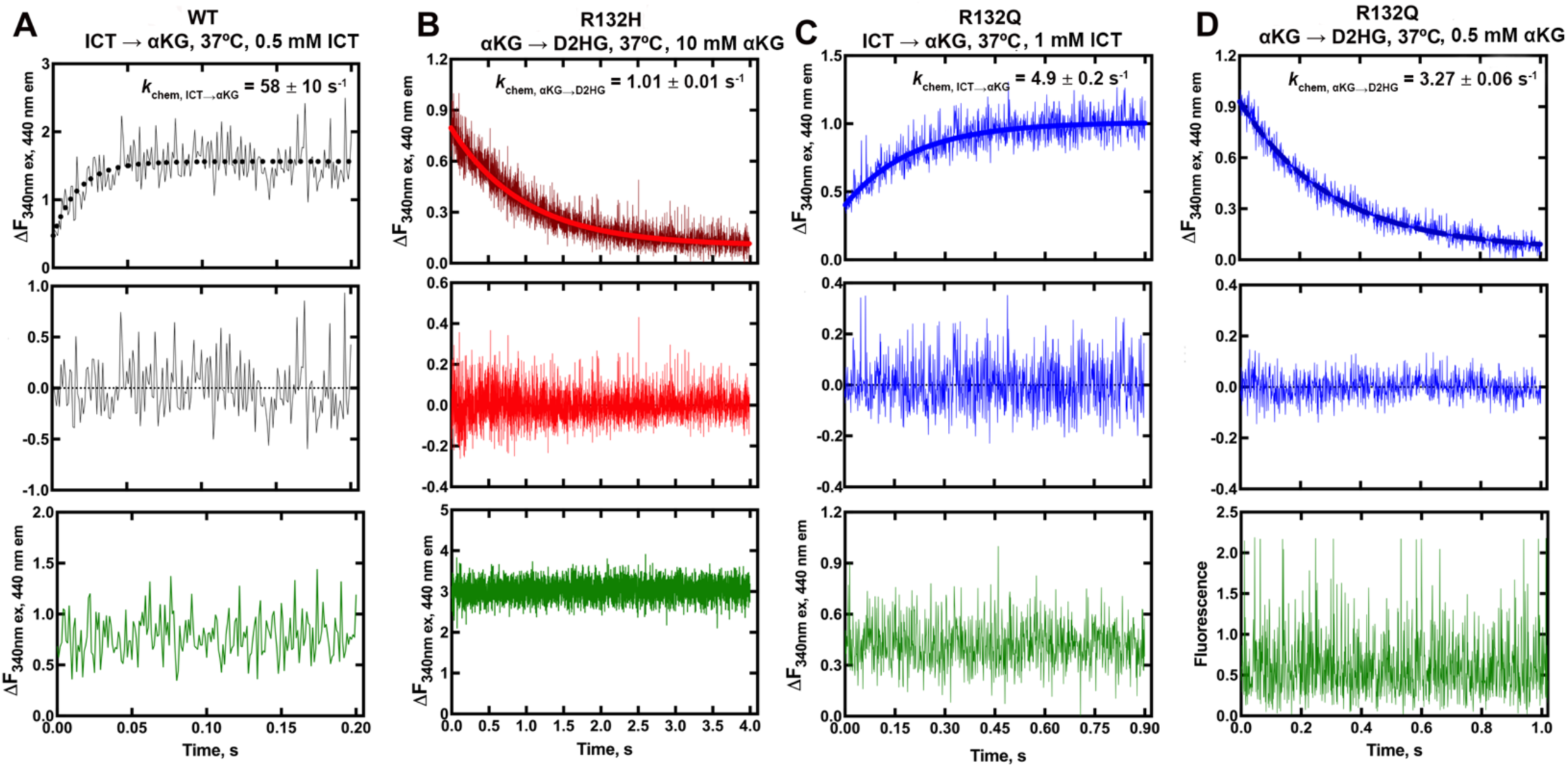
Pre-steady-state single-turnover kinetic features of IDH1 WT, R132H, and R132Q catalysis. NADPH formation in the conventional reaction and consumption in the neomorphic reaction was monitored over the course of a single turnover (top plot) and compared with a control experiment lacking enzyme (bottom plot, in green). Residuals (middle plot) were obtained to assess goodness of a single exponential equation fit. A) IDH1 WT, conventional reaction. B) IDH1 R132H, neomorphic reaction. C) IDH1 R132Q, conventional reaction. D) IDH1 R132Q, neomorphic reaction.

We were unable to capture rates of conformational change when monitoring intrinsic protein fluorescence. However, we measured rates of NADPH binding to IDH1 WT, R132H, and R132Q using enzyme that was stripped of cofactor ^14^ (Extended Data Fig. 3). We found that all three IDH1 proteins displayed single-step binding events, with an NADPH binding on rate (*k*_on_) for IDH1 WT that was ∼2-fold faster than that calculated for IDH1 R132Q, while *k*_on_ rates for IDH1 R132H were profoundly slower. We also used isothermal titration calorimetry (ITC) to measure equilibrium binding affinity of NADPH for IDH1 (Supplementary Fig. 1). We found that both mutants exhibited a 5-fold decrease in *K*_d_ compared to IDH1 WT, suggesting that a slower *k*_off_ rate drove the improved affinity for NADPH observed for R132H despite the slow *k*_on_ rate. Taken together, these kinetic data further supported the finding that when compared to R132H, IDH1 R132Q has a lower barrier to adopting the closed, active conformation that is driven by substrate and metal binding.

### IDH1 R132Q has a less solvent accessible active site pocket that is more catalytically primed

To illuminate possible mechanisms behind the time-resolved changes exhibited by IDH1 R132Q versus those in WT and R132H, we first used hydrogen/deuterium exchange-mass spectrometry (HDX-MS) analysis. We probed solvent accessibility as indicated by deuterium uptake in the binary IDH1:NADP(H) form, as WT and mutant IDH1 are known to copurify bound to NADP(H) ^26,27^. We also measured deuterium uptake upon the addition of substrate (ternary complex, IDH1:NADP(H):ICT/αKG), or upon the addition of substrate and Ca^2+^ (quaternary complex, IDH1:NADP(H):ICT/αKG:Ca^2+^). By far the most substantial change in deuterium uptake for WT, R132H, and R132Q occurred in the quaternary form, indicative of closed, catalytically competent conformations among all enzyme species (Extended Data Fig. 4). This is consistent with previous findings that both substrate (ICT, but also presumably αKG in the neomorphic reaction) and divalent metal binding are required to drive IDH1 into its fully closed, active conformation ^25–28^. Deuterium uptake generally showed the following trend: R132H:NADPH:αKG:Ca^2+^ >> WT:NADP^+^:ICT:Ca^2+^ > R132Q:NADPH:ICT:Ca^2+^ > R132Q:NADPH:αKG:Ca^2+^ (Fig. 2, Extended Data Fig. 5), with R132Q appearing to have an overall less structurally dynamic, more closed conformation compared to R132H.

**Fig. 2.**
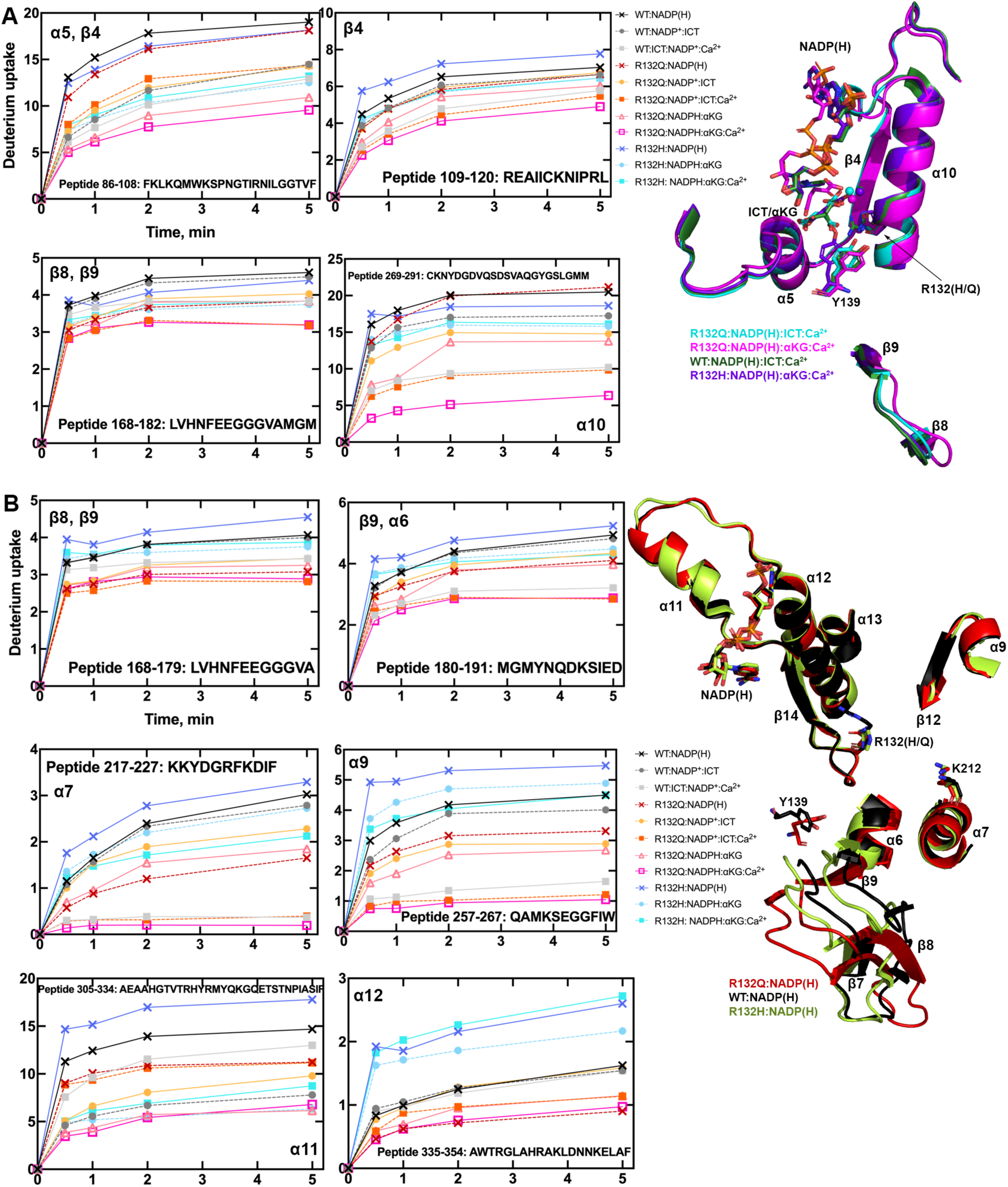
IDH1 R132Q has lower deuterium uptake than R132H in both binary and quaternary complexes. A) Plots of deuterium uptake encompassing residues 86-123, 168-209, and 260-291 (left), with the structural features of these residues shown in cartoon (right) for IDH1 R132Q, WT ^13^, and R132H ^14^. B) Plots of deuterium uptake for residues 144-191, 217-227, 257-267, and 295-354 (left), with the structural features of IDH1 R132Q, WT ^13^, and R132H ^29^ encompassing these regions are shown in cartoon (right).

Since our kinetic studies suggested IDH1 R132Q had a lower barrier to achieve the closed conformation compared to R132H, we hypothesized that the binary R132Q:NADP(H) would be in a more quaternary-like state. To test this, we compared deuterium uptake among the binary states, predicting that the R132Q:NADP(H) complex would experience less deuterium uptake than R132H:NADP(H). Unsurprisingly, in general the IDH1:NADP(H) forms of all three proteins had high deuterium uptake, particularly in the substrate binding pocket, clasp domain, and dimer interface (Fig. 2, Extended Data Figs. 4, 5). As predicted, R132Q:NADP(H) and WT:NADP(H) had the least deuterium uptake overall, while R132H:NADP(H) exhibited, by far, the most uptake. As this suggested that NADP(H)-bound R132Q had a more closed/less mobile conformation compared to R132H, we wondered if the temporal features of our HDX-MS data suggested a faster closing upon substrate binding for the R132Q mutant. This would provide one mechanism of the improved catalytic efficiency shown by IDH1 R132Q relative to R132H in both the conventional and neomorphic reactions. To address this, we inspected peptides that included residues within 4 Å of the bound NADP(H) and ICT/αKG substrates to determine if deuterium uptake equilibrium was reached faster in IDH1 R132Q versus R132H, consistent with a primed ground state that reached a closed conformation more easily. Interestingly, the peptides that contained active site residues from the adjacent monomer (i.e., chain B residues contributing to the chain A active site) showed a faster approach to deuterium uptake equilibration for IDH1 R132Q and WT compared to R132H (Extended Data Fig. 6). Specifically, peptides 210-216, 240-253, and 257-267 all showed IDH1 R132Q reaching an equilibrium state faster than R132H. This is supportive of a model where the ground state of R132Q is a more closed conformation that follows a simpler path to a catalytically competent state compared to R132H.

Seeking to pair the dynamic, intermediate-resolution HDX-MS data with static, high-resolution X-ray crystal structures, we report here six new crystallographic models representing the first structures of IDH1 R132Q: binary IDH1 R132Q bound to NADP(H) (R132Q:NADP(H), PDB ID 8VHC and 8VH9); R132Q bound to conventional reaction substrates (R132Q:NADP(H):ICT:Ca^2+^, PDB ID 8VHD); R132Q bound to neomorphic reaction substrates (R132Q:NADP(H):αKG:Ca^2+^, PDB ID 8VHB and 8VHA), and R132Q bound to a NADP-TCEP adduct (R132Q:NADP-TCEP:Ca^2+^, PDB ID 8VHE). These structures facilitated comparisons with previously solved IDH1 WT ^13^ and R132H structures ^14,29^, including among binary and ICT- and αKG-bound models.

Binary structures of IDH1 R132Q were valuable to help us understand how the active site of cofactor-bound R132Q compared to R132H. While R132Q:NADP(H) showed no major global structural alterations upon alignment with previously solved structures of WT:NADP(H) ^13^ and R132H:NADP(H) ^29^, many local shifts were observed (Fig. 3). Unsurprisingly, NADP(H)-bound R132Q had the typical overall open, inactive conformation seen in WT and R132H, with a larger active site cleft and smaller back cleft relative to the quaternary complexes (Supplementary Table 1). Interestingly, these distances in the binary IDH1 R132Q structure more closely resembled binary WT than R132H, supportive of a more closed, catalytically competent ground state for R132Q. However, IDH1 R132Q exhibited notable differences compared to WT and R132H. In particular, the clasp domain and helices proximal to the substrate and cofactor binding site were shifted, with the α1, α2, α4, α5, and α11 helices adjusted upwards and inwards in R132Q versus WT and R132H binary complexes, resulting in a similar shift of the NADP(H) molecule itself (Fig. 3B). Importantly, this inward shifting of the α1 helix is a feature of closed, catalytically competent IDH1 conformations. Notably, R132Q also contained longer, more intact β strands in the clasp domain, which plays a major role in maintaining the dimer, compared to both WT and R132H (Fig. 3). The fully intact β7 and β8 strands in R132Q were reminiscent of quaternary, fully substrate-bound forms of IDH1 WT and R132Q (*vide infra*). Consistent with such stable secondary structure, peptides in the β8 strand of R132Q:NADP(H) had lower deuterium uptake than WT:NADP(H) and R132H:NADP(H) (Fig. 2, Extended Data Fig. 5). IDH1 R132Q also maintained an extensive hydrogen bonding network enveloping the NADP(H) molecule; this network was far less robust in R132H (Extended Data Fig. 7). Together, both dynamic and static structural data suggest that the IDH1 R132Q active site pocket and surrounding features have greater rigidity and more defined structural features typical of fully-substrate-bound forms of IDH1, suggesting a more catalytically primed state for R132Q:NADP(H) compared to R132H:NADP(H).

**Fig. 3.**
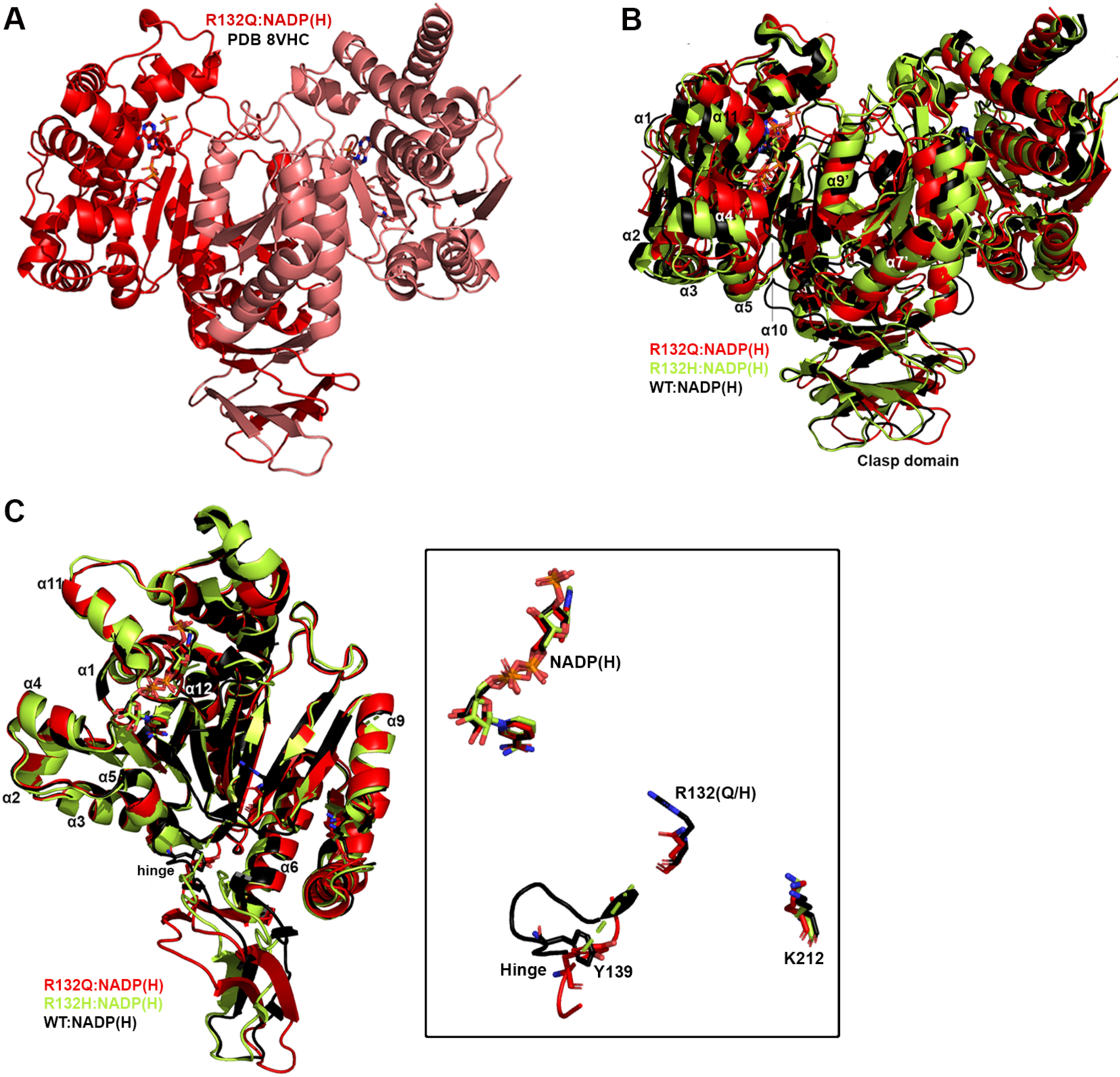
Crystal structure of NADP(H)-bound IDH1 R132Q. A) The binary R132Q:NADP(H) complex is shown with each monomer highlighted using a slight color change. B) Dimer-based alignments of R132Q:NADP(H), WT:NADP(H) ^13^, and R132H:NADP(H) ^29^. C) Monomer-based alignments of R132Q:NADP(H), WT:NADP(H) ^13^, and R132H:NADP(H) ^29^. The inset features catalytic residues Y139 and K212 (though the latter residue drives catalysis in the monomer not shown), residue R132(H/Q), and the cofactor.

### Unlike R132H, ICT-bound IDH1 R132Q is in a closed, catalytically competent conformation

Here, we also report the first ICT-bound quaternary structure of IDH1 R132Q (R132Q:NADP(H):ICT:Ca^2+^). Upon alignment of this structure with WT:NADP(H):ICT:Ca^2+^ ^13^ (Fig. 4), there was obvious overlap in both global features and active site details. ICT-bound IDH1 R132Q also aligned well with R132H bound to its preferred substrate, αKG (R132H:NADP(H):αKG:Ca^2+^) ^14^. Like ICT-bound WT and αKG-bound R132H structures, ICT-bound IDH1 R132Q adopted a catalytically competent, closed conformation, with ICT maintaining many of the same polar interactions with the protein and divalent ion as observed with IDH1 WT. This is supportive of our kinetic data showing R132Q’s preservation of the conventional activity.

**Figure 4.**
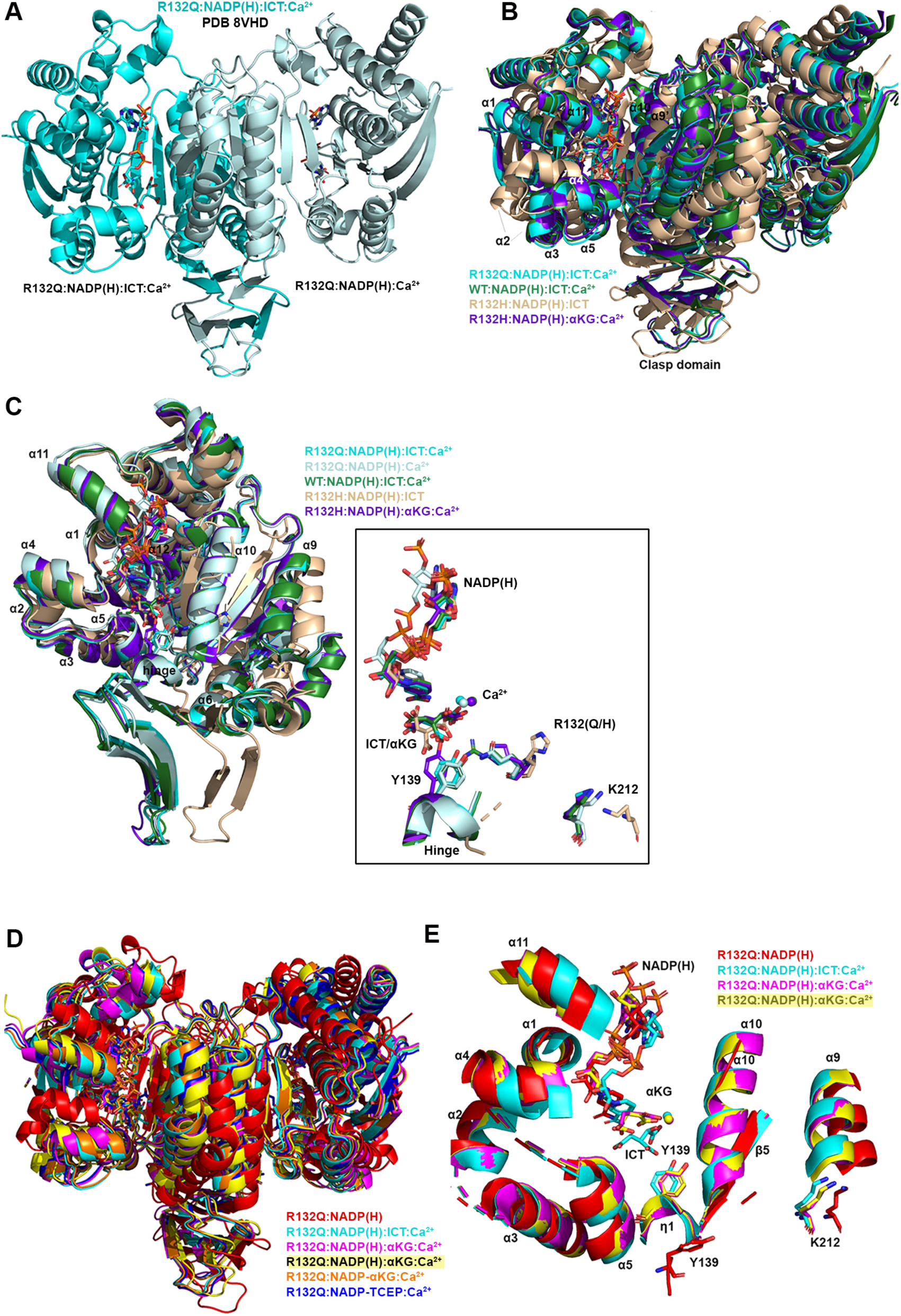
Crystal structure of IDH1 R132Q bound to ICT, NADP(H) and Ca^2+^ that mimics the catalytic Mg^2+^. A) The R132Q:NADP(H):ICT:Ca^2+^ complex is shown with each monomer highlighted using a slight color change. B) Dimer-based alignments of the R132Q:NADP(H):ICT:Ca^2+^/R132Q:NADP(H):Ca^2+^ dimer (cyan); R132H:NADP(H):ICT ^29^ (wheat); R132H:NADP(H):αKG:Ca^2+^ ^14^ (dark purple); and WT:NADP(H):ICT:Ca^2+^ ^13^ (dark green). C) Monomer-based alignments of the R132Q:NADP(H):ICT:Ca^2+^/R132Q:NADP(H):Ca^2+^ dimer (dark and light cyan) with structures described in (B). For clarity, only the catalytic residues, residue R132X, cofactor, substrates, Ca^2+^ and hinge are shown in the inset. D) Dimer-based alignments of R132Q:NADP(H), R132Q:NADP(H):ICT:Ca^2+^, and R132Q:NADP(H):αKG:Ca^2+^. E) Monomer-based alignment of the IDH1 R132Q structures described in (D) with substrates, catalytic residues, and nearby secondary structure features highlighted.

Though alignment of ICT-bound WT and R132Q was strikingly similar, the notable 180-fold decrease in catalytic efficiency suggested that maintaining hydrogen bonding features and active site structuring was not sufficient for robust conventional activity in R132Q. Interestingly, ICT was observed only in one monomer of the R132Q quaternary complex, resulting in a shift of the α11 helix and the NADP(H) molecule upward and outward in the ICT-absent R132Q monomer (Fig. 4), reminiscent of the WT:NADP(H) binary structure (Fig. 3). This lack of active site saturation suggested a lower affinity toward ICT for IDH1 R132Q versus WT. Though *K*_m_ values are not affinity measurements, it is noteworthy that there was a >300-fold increase in *K*_m_ when comparing R132Q to WT. To address differences in binding affinity, we again turned to ITC experiments. While ICT binding affinity for IDH1 R132H was below the limit of detection, we found that R132Q exhibited a ∼170-fold decrease in ICT affinity compared to WT (Supplementary Fig. 1). Further, evidence of ICT binding was observed for R132Q, but not R132H. Structural studies again provided a possible mechanism; in contrast to the closed, catalytically competent conformation of ICT-bound R132Q, a previously solved ternary R132H:NADP(H):ICT ^29^ structure revealed quasi-open monomers that had α4 and α11 helices shifted upwards and outwards from the dimer interface and an unraveled α10 helix (Fig. 4B, 4C), regions we and others have shown to be highly flexible ^13,29–31^. Notably, ICT was found in a posited pre-binding site that was shifted to the left of its catalytically-competent position ^29^. This resulted in limited polar interactions by ICT to R132H ^29^ in contrast to ICT’s extensive polar contacts to R132Q, including hydrogen bonding to catalytic residue Y139 that indicated a catalytically-ready binding conformation (Extended Data Fig. 7). As further evidence that ICT-bound R132H was ill-prepared for catalysis, its catalytic residues swung away from the active site, akin to the positioning found in binary, catalytically incompetent IDH1 structures (Fig. 4C). Though this IDH1 R132H structure did not include a divalent metal that may be required for full closure ^29^, it is nonetheless unsurprising that IDH1 R132H, in contrast to IDH1 R132Q, is essentially unable to convert ICT to αKG.

### IDH1 R132Q αKG-bound form is semi-closed, with an αKG binding pocket that is unique from R132H

Since IDH1 R132Q uniquely maintains both normal and neomorphic catalytic abilities, we asked how the binding conformations for ICT, the conventional reaction substrate, and αKG, the neomorphic reaction substrate, compared. Here, we report two αKG-containing IDH1 R132Q quaternary structures (R132Q:NADP(H):αKG:Ca^2+^). These co-crystallization experiments led to a variety of complexes, with monomer asymmetry still observed (Fig. 5). One structure had αKG bound in one monomer, and a covalent NADP-αKG adduct in the other (Fig. 5A, Supplementary Fig. 2). Cleft measurements in both monomers indicated a slightly more open conformation when compared to the closed quaternary R132Q (ICT-bound), WT (ICT-bound) and R132H (αKG-bound) structures, with the α11 helix shifted out away slightly from the substrate binding pocket (Fig. 5E). As a result, the NADP(H) itself shifted outwards compared to the ICT-bound R132Q structure, resulting in a semi-closed conformation (Supplementary Table 1).

**Figure 5.**
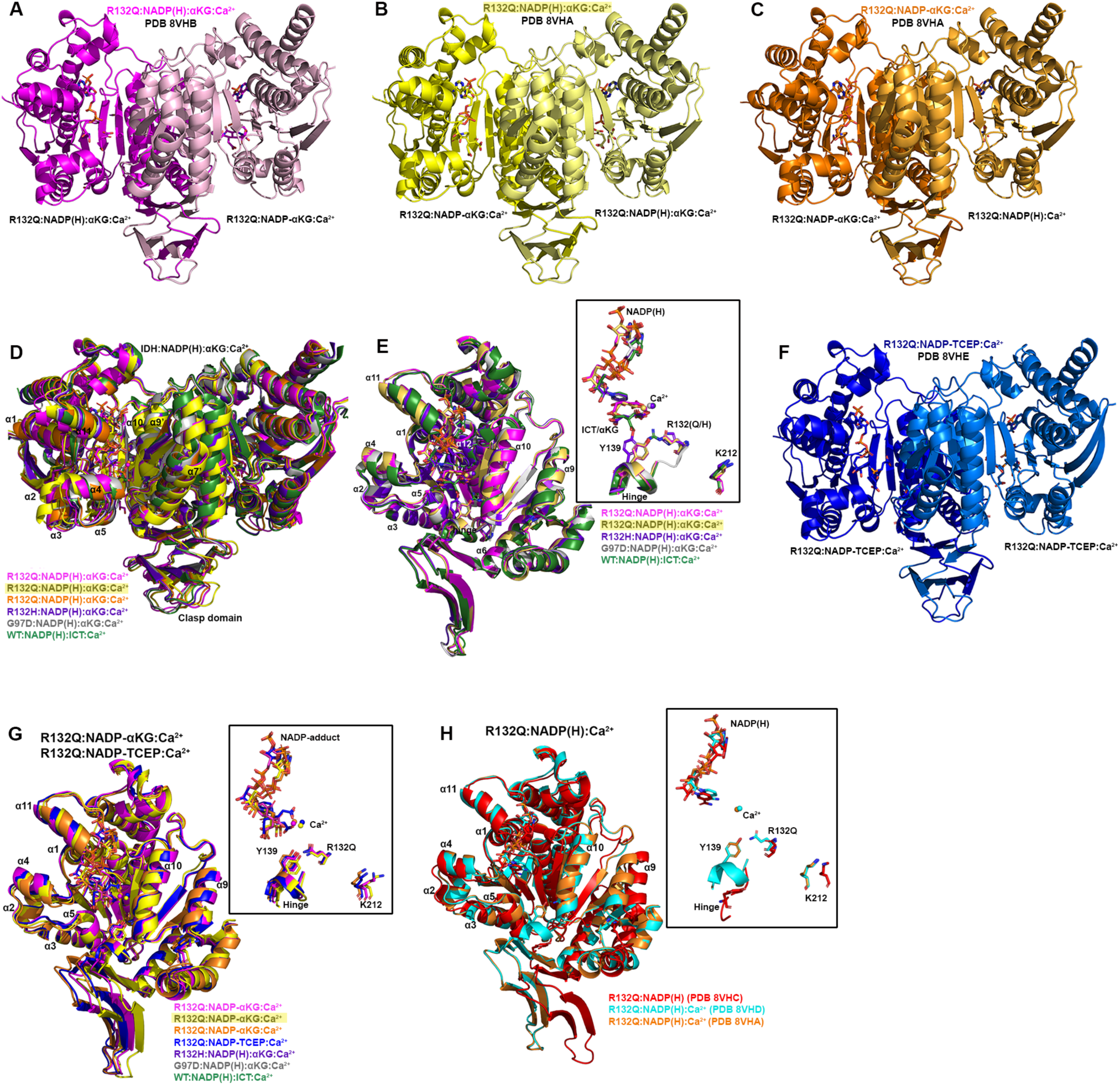
Crystal structure of IDH1 R132Q bound to αKG and NADP-adducts. In (A-C) and (F), each R132Q monomer is highlighted using a slight change in color, with a description of the ligands listed below each monomer. A) R132Q:NADP(H):αKG:Ca^2+^/R132Q:NADP-αKG:Ca^2+^ dimer. B) R132Q:NADP-αKG:Ca^2+^/ R132Q:NADP(H):αKG:Ca^2+^ dimer 1. C) R132Q:NADP-αKG:Ca^2+^/R132Q:NADP(H):Ca^2+^ dimer 2. D) Dimer-based alignments of R123Q αKG-bound structures with R132H:NADP(H):αKG:Ca^2+^ ^14^, G97D:NADP(H):αKG:Ca^2+^ ^14^, and WT:NADP(H):ICT:Ca^2+^ ^13^. E) Monomer-based alignment of αKG-containing R132Q monomers with structures described in (D). F) R132Q:NADP-TCEP:Ca^2+^/R132Q:NADP-TCEP:Ca^2+^ dimer. G) Alignment of adduct-containing R132Q monomers. H) Alignment of the non-substrate containing R132Q monomers.

The second αKG-bound structure had unique features among two dimers in the crystallographic asymmetric unit. One catalytic dimer contained one NADP-αKG adduct and one αKG molecule (Fig. 5B), and again appeared as an intermediate between the R132Q:NADP(H) and the R132Q:NADP(H):ICT:Ca^2+^ structures (Supplementary Table 1). A second dimer contained an NADP-αKG adduct in one monomer, while no αKG-containing molecule was observed in the adjacent monomer (Fig. 5C). This dimer was in a more closed, catalytically competent conformation, reminiscent of the fully closed WT quaternary structure (Supplementary Table 1). The Ca^2+^ ion clearly led to extensive restructuring, as the R132Q:NADP(H):Ca^2+^ monomer aligned relatively poorly with the R132Q:NADP(H) complex despite the only difference being the presence of the metal ion (Fig. 5H). Thus, closing of IDH1 R132Q to the αKG-bound form may be driven just as much by metal binding as by substrate binding. This finding was recapitulated by the overall decrease seen in deuterium uptake upon treatment of substrate-bound R132Q with Ca^2+^ (Extended Data Fig. 4). Overall, we were able to capture snapshots of stable conformations of αKG binding ranging from semi-closed (αKG-bound) to essentially fully closed (NADP-αKG adduct-bound).

Closed conformations are seen for WT ^13^ and R132H ^14^ when bound with their preferred substrates (ICT and αKG, respectively). As αKG-bound R132Q was often not as fully closed as the ICT-bound form, we wondered how αKG-bound R132Q compared to these WT and R132H closed conformations. In alignments of R132Q:NADP(H):αKG:Ca^2+^ with quaternary WT and R132H structures (Fig. 5D, 5E), the catalytic residue Y139 in R132Q was shifted away from the αKG molecule, with this molecule making fewer hydrogen bond contacts within the R132Q active site compared to R132H (Extended Data Fig. 7). In R132Q, the αKG binding site was shifted upwards towards NADP(H) and away from the substrate binding sites seen in the ICT-bound WT and αKG-bound R132H structures. This shift might be facilitated by one surprising feature of all non-αKG-containing R132Q monomers -- the nicotinamide ring could not be reliably modeled due to missing electron density (Fig. 5). This suggests that when αKG was absent (such as in the R132Q:NADPH:Ca^2+^ monomer that dimerized with the NADPH-αKG adduct) or, more unexpectedly, even when αKG was bound (R132Q:NADPH:αKG:Ca^2+^ monomers), this portion of NADP(H) was more dynamic in the active site. Overall, the αKG-containing R132Q structures either did not appear in a catalytically-ready form, or the enzymatic mechanism may rely more heavily on different amino acids used in the conventional reaction.

### ICT-bound and αKG-bound IDH1 R132Q show unique static and dynamic structural features

The α10 regulatory segment undergoes notable restructuring upon substrate binding, with this segment forming a helix in both the ICT- and αKG-bound quaternary forms of R132Q, just like in ICT-bound WT and αKG-bound R132H (Figs. 4, 5). However, our HDX-MS experiments captured more subtle differences in R132Q that depended on the substrate that was bound. The α10 regulatory segment and the nearby α9 helix were much more protected from proton exchange in both αKG and αKG + Ca^2+^ conditions in R132Q than in the ICT and ICT + Ca^2+^ conditions (Figs. 2, 6). Beyond its proximity to the regulatory segment, the α9 helix has an additional role in active site remodeling in that it helps form a “seatbelt” that envelopes the NADP(H) cofactor in many aldo-keto reductase enzymes (reviewed in ^32^). This seatbelt was observed in the WT:NADP(H):ICT:Ca^2+^ quaternary structure, with residue R314 in α11 helix shifted inward to form polar contacts with D253’ and Q256’ in α9 of the adjacent monomer and with a water molecule (Fig. 7). The absence of the seatbelt was not limited to binary R132Q, R132H, and WT structures; no seatbelt was observed in the ternary ICT-bound or, more surprisingly, in the closed, quaternary αKG-bound R132H structures ^14,29^. As no αKG-bound IDH1 WT structure is available at this time, we compared a structure of a non-R132 mutant, G97D, which generates D2HG but exhibits a high degree of structural similarities with IDH1 WT ^14^. The αKG-bound form of G97D also did not show a seatbelt conformation, suggesting this is a unique feature of ICT-bound, fully closed structures.

**Fig. 6.**
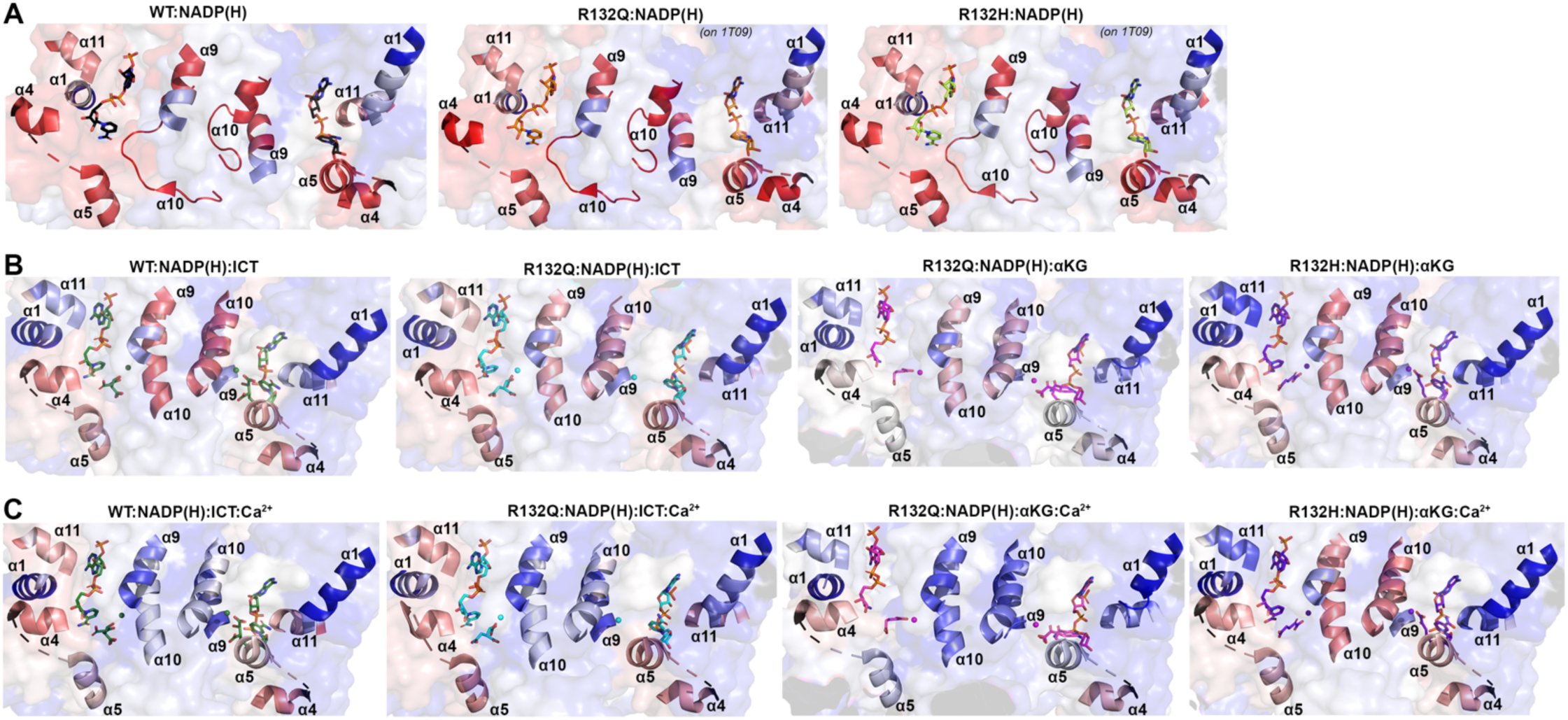
Deuterium uptake by IDH1 WT, R132Q, and R132H in helices bounding the substrate binding pocket. Deuterium uptake is shown as a gradient from red (high uptake) to blue (low uptake). A) Deuterium uptake by IDH1 WT, R132Q, and R132H upon no ligand treatment. These HDX-MS data were overlaid on NADP(H)-only bound forms of WT ^13^ in all three cases, as the αKG helix was disordered in the NADP(H)-only bound forms of IDH1 R132Q and R132H ^29^. B) Deuterium uptake by WT and R132Q upon treatment with NADP^+^ and ICT, and by IDH1 R132Q and R132H upon treatment with NADPH and αKG. These HDX-MS data were overlaid on WT:NADP(H):ICT:Ca^2+^ ^13^, R132Q:NADP(H):ICT:Ca^2+^ and R132Q:NADP(H):αKG:Ca^2+^, or R132H:NADP(H): αKG:Ca^2+^ ^14^. C) Deuterium uptake by IDH1 WT and R132Q upon treatment with NADP^+^, ICT, and Ca^2+^, and by IDH1 R132Q and R132H upon treatment with NADPH, αKG, and Ca^2+^. These HDX-MS data were overlaid on the structures described in (B).

**Fig. 7.**
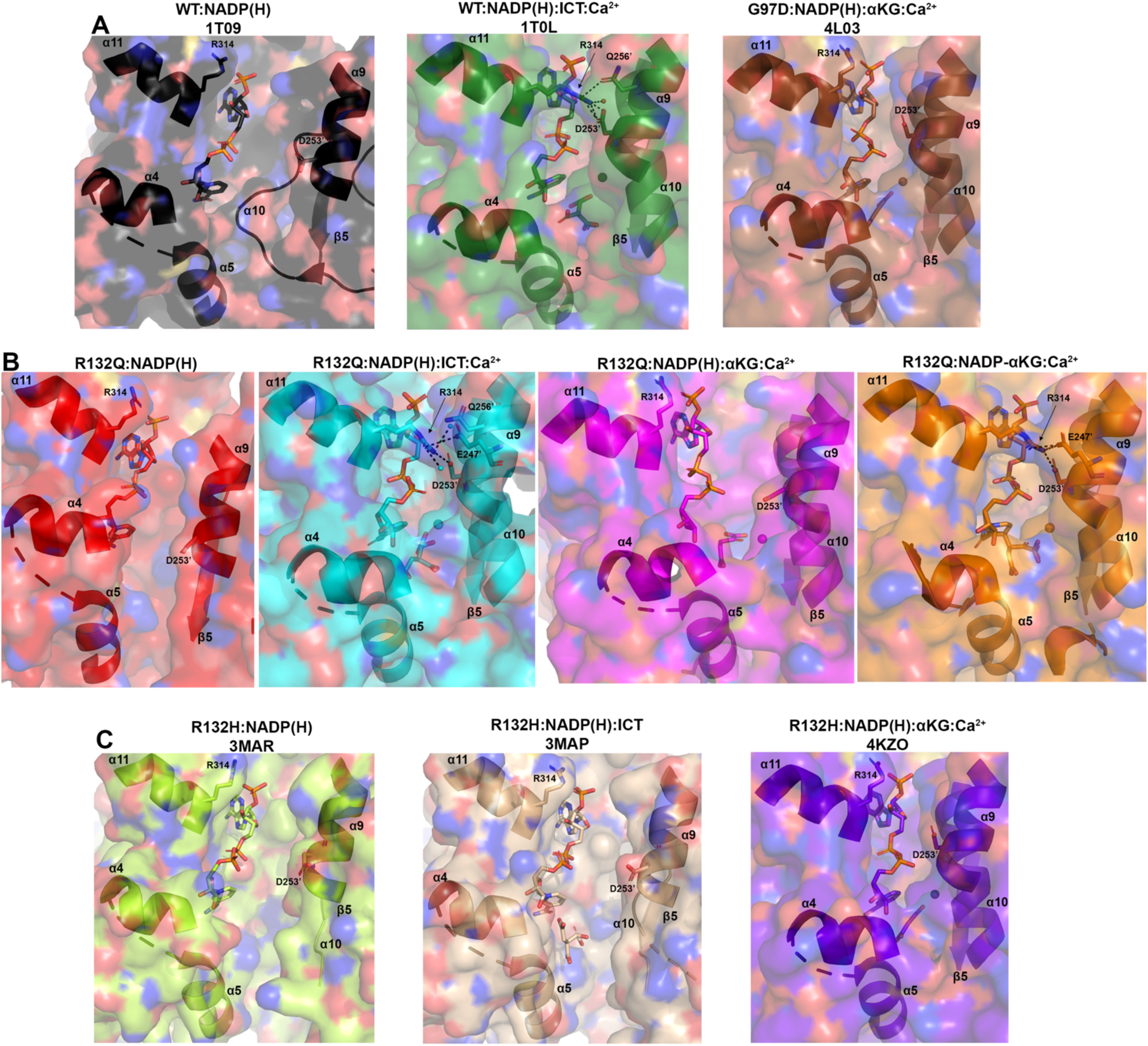
Hydrogen bond network facilitates a “seatbelt” that overlays NADP(H) in only some quaternary structures of IDH1. A) Unlike the binary structure of IDH1 WT ^13^ and quaternary structure of G97D:NADP(H):αKG:Ca^2+^ ^14^, the quaternary IDH1 WT complex ^13^ forms a seatbelt over the NADP(H). B) Binary R132Q:NADP(H) and quaternary R132Q:NADP(H):αKG:Ca^2+^ structures do not form a seatbelt, while R132Q:NADP(H):ICT:Ca^2+^ and the most closed conformation of R132Q:NADP-αKG:Ca^2+^ for a seatbelt. C) No seatbelt is formed in the binary R132H:NADP(H), ternary R132H:NADP(H):ICT, or quaternary R132H:NADP(H):αKG:Ca^2+^ structures of IDH1 R132H ^14,29^.

IDH1 R132Q behaved like WT when binding the conventional reaction substrate (ICT), with a seatbelt forming over the cofactor since residue R314 was in position to contact Q256’, D253’, and, unique to this protein, E247’ in β11 of the adjacent monomer, as well as a water molecule (Fig. 7). However, R132Q behaved more like R132H when binding the neomorphic substrate, with αKG-bound monomers showing residue R314 swung away from the α9’ helix, precluding the necessary polar contacts. Interestingly, the closed R132Q:NADP-αKG:Ca^2+^/R132Q:NADP(H):Ca^2+^ dimer (Fig. 5C) had an intact seatbelt over the NADP-αKG adduct (Fig. 7B), suggesting that a fully closed conformation of αKG-bound IDH1 R132Q is possible if the nicotinamide ring of NADP(H) is stabilized in some way, such as via adduct formation. Interestingly, HDX-MS dynamics showed that seatbelt formation was associated with an increase in deuterium uptake, with the α11 helix, which contains the seatbelt-forming R314 residue, more protected in the αKG-bound R132Q and R132H (seatbelt-lacking) complexes relative to the ICT-bound WT and R132Q (seatbelt-forming) complexes (Fig. 6). Overall, multiple conformations are possible with αKG-containing R132Q structures, including those associated with fully closed forms.

### IDH1 R132Q accommodates multiple NADP-containing adducts that may provide clues to transition-state features

In addition to the NADP-αKG adduct, we also encountered an NADP-tris(2-carboxyethyl)phosphine (NADP-TCEP) adduct when attempting to crystallize ICT-bound R132Q (Fig 5F, Supplementary Fig 2). There appeared to be some catalytic relevance of these adducts in that the TCEP and αKG carboxylates in the adducts helped coordinate the Ca^2+^ and maintained many hydrogen bonds in their respective active sites, though the metal ion was slightly shifted to accommodate the adducts (Fig. 5, Supplementary Fig. 4). All TCEP and αKG adducts appeared as hybrids between the semi-closed αKG-bound R132Q complex and fully closed ICT-bound R132Q complex (Supplementary Table 1). In general, One NADP-αKG adduct-containing monomer (Fig. 5C) aligned well to the fully closed ICT-bound R132Q structure in all regions except the clasp domain, where the adducted monomer was shifted towards the dimer interface and the β9 strand was more intact (Fig. 5G). As further evidence of its fully closed conformation, this NADP-αKG adduct-containing monomer also had an intact seatbelt feature (Fig. 7B).

To better understand how these adducts were forming, we performed density functional theory (DFT) calculations for model NADP-TCEP and NADP-αKG adducts (Supplementary Tables 2, 3), which suggested that adduct formation would fail to occur if not for the constraining environment of the crystal structure. We considered an alternative possibility that the IDH1 R132Q active site itself favored adduct formation and binding. If the NADP-TCEP adduct could form in the active site of R132Q, it would act as a competitive inhibitor. Thus, we treated R132Q with varying concentrations of three reducing agents (TCEP, dithiothreitol (DTT), and β-mercaptoethanol (BME)) to determine the effects of catalysis of the conventional reaction (Extended Data Fig. 8, Supplementary Table 4). Dose-dependent inhibition of R132Q catalysis was profound with TCEP, while DTT and DME had minimal effects. More modest, though notable effects on catalysis were also observed when challenging IDH1 WT with the highest concentration of TCEP tested (10 mM) (Supplementary Fig. 3). As NADP-DTT adducts have been previously reported ^33^, our discovery that DTT did not inhibit R132Q may help support a model where the enzyme supports adduct formation. While DTT has some similar structural features compared to ICT, it does not recapitulate the carboxylic acid features that TCEP and αKG provide in their NADP-containing adducts. Together, these results strongly support the hypothesis that adduct formation occurs outside of the non-physiologically-relevant crystal packing environment, with the adducts mimicking αKG binding, ICT binding, or transition between the two.

As these adduct-containing structures showed hybrid binding features of αKG and ICT, we wondered if transition state features could be extrapolated. Here, the nicotinamide ring of the adduct lent an interesting clue. Calculations suggest that the nicotinamide ring is likely to be planar in the oxidized form ^34,35^. During NADP^+^ activation for hydride transfer, the enzyme is predicted to distort the nicotinamide ring to form a highly puckered transition state as a partial positive charge on C4N develops ^34–36^ (Supplementary Fig. 5). NAD(P)-adducts with reducing agents have been reported previously, including with TCEP ^37^ and also with DTT ^33^, and were often found to have a more puckered nicotinamide ring, reminiscent of a transition state. Here, unlike the planar ring observed in our non-adducted forms of NADP(H) (R132Q:NADP(H):ICT:Ca^2+^), both the αKG- and TCEP-containing NADP adducts showed a more puckered nicotinamide ring (Supplementary Fig. 4, Supplementary Table 3), suggestive of a transition-state-like conformation.

In summary, we highlight discrete catalytic and structural features among two tumor-relevant IDH1 mutants, with the IDH1 R132Q mutant serving as an invaluable tool to probe the journey through substrate turnover of two reactions that typically cannot be performed by the same enzyme. Together, our kinetics experiments and static and dynamic structural data suggests that substrate binding and conformational changes associated with the conventional and the neomorphic reactions have unique paths through turnover that can be described in terms of differences in substrate affinity, substrate binding site location, solvent accessibility, and propensity for conformational activation and active site remodeling (summarized in Fig. 8). IDH1 R132Q’s accommodation of catalytically-relevant adducts, perhaps due to its active site appearing better optimized for catalysis compared to R132H, illuminate snapshots of substrate and substrate analogs in varying degrees of catalytic readiness.

**Fig. 8.**
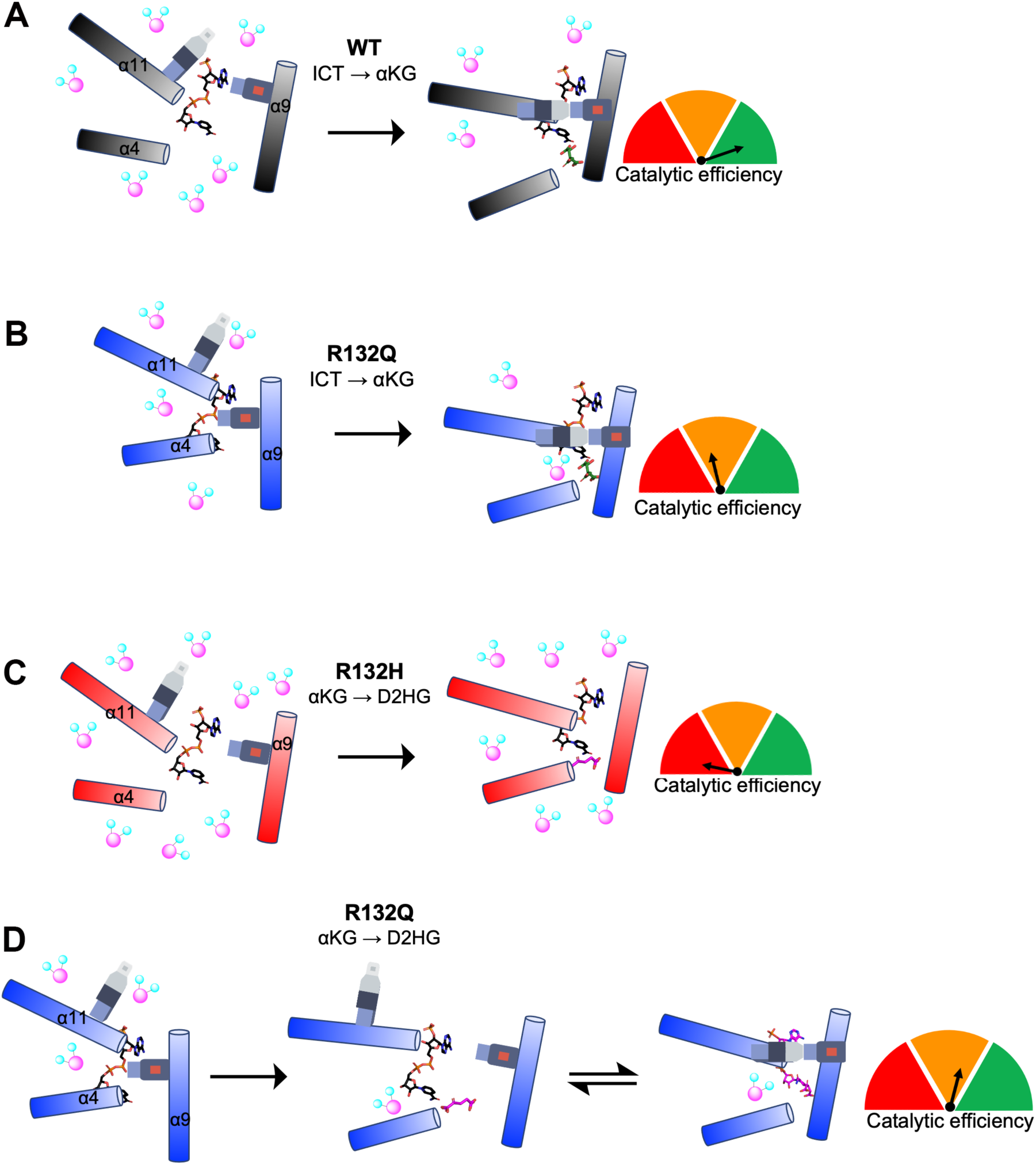
Conformations and solvent accessibility of IDH1 WT, R132Q, and R132H upon substrate binding. Helices displaying profound differences in alignment of the three forms of IDH1 are highlighted. The seatbelt feature is indicated on the α11 and α9 helices. A) Binary WT:NADP(H) ^13^ collapses to a closed conformation upon ICT binding, though moderate levels of deuterium exchange are still permitted. B) Binary R132Q:NADP(H) collapses to a closed conformation upon ICT binding, showing improved catalytic efficiency for the conventional reaction and lower deuterium uptake compared to R132H. However, catalytic activity is much lower compared to WT. C) Binary R132H:NADP(H) ^29^ collapses to a fully closed conformation only upon αKG binding ^14^, but a seatbelt is not formed and deuterium uptake remains high. D) Binary R132Q:NADP(H) forms semi-closed and closed conformations upon binding αKG and NADP-αKG, respectively, with a seatbelt successfully formed in the closed state. The αKG binding site was shifted away from the α9 helix, though catalytic activity was much higher than that seen in R132H.

## Discussion

Steady-state and pre-steady-state kinetic, HDX-MS, and X-ray crystallography experiments revealed fundamental differences in the molecular mechanisms of catalysis by WT and tumor-relevant IDH1 mutants (Fig. 8). It is unsurprising that IDH1 WT is far more efficient at catalyzing the conventional reaction than R132Q and R132H since R132 coordinates the C3 carboxylate of isocitrate ^3,13^. As neither mutant can directly participate in this coordination, we asked why the conventional reaction was more efficient in R132Q than R132H. We found that R132Q employed a unique active site water that mitigated the loss of hydrogen bonding to ICT resulting from the R to Q mutation by imperfectly mimicking the polar interactions with the substrate normally afforded by R132 (Extended Data Fig. 9). Despite the shifting of the αKG binding site, we noticed a similar compensatory mechanism in our αKG-bound R132Q structure, with a water molecule recapitulating the polar interactions normally made by residue R132. Here, however, the water molecule did not appear to hydrogen bond with the substrate. Instead, a second water molecule was found at the same location as the Ca^2+^ ion in the quaternary ICT-bound WT IDH1 structure (Extended Data Fig. 9), which presumably helped stabilize the αKG substrate in R132Q. We have previously reported the importance of water molecules in facilitating mutant IDH1 inhibition ^30^, and this current work highlights the importance of water in substrate binding by providing a possible mechanism by which R132Q is more catalytically efficient compared to R132H.

In addition to affecting catalysis, the α10 regulatory segment may also serve as a selectivity filter for mutant IDH1 inhibitor binding ^38^. We have shown previously that IDH1 R132Q binds poorly to selective mutant IDH1 inhibitors, with IC50 profiles consistent with IDH1 WT rather than R132H ^22^. We had predicted that a much more stable α10 regulatory segment in R132Q:NADP(H) drove this resistance. Here, we found that while this unfolded loop indeed had stronger electron density compared to R132H:NADP(H) ^29^, it still appeared less stable than the partially folded features of WT:NADP(H) ^13^. Instead, we now believe that the more activated, quaternary-like state of the binary R132Q:NADP(H) complex helps drive inhibitor resistance. In the binary R132Q complex, regions including the α11 and α4 helices were shifted inwards and the protein experienced less deuterium uptake (Fig. 8). Using compound 24 as a prototypical selective mutant IDH1 inhibitor (6O2Y ^39^), it did not appear that the small increase in the stability of the α10 regulatory segment in IDH1 R132Q would have much of an effect on inhibitor binding (Extended Data Fig. 10). Instead, our alignments showed residues 111-121 in the inhibitor binding pocket, which form a loop between the β4 and β5 strands, likely have a larger role in the loss of affinity towards inhibitors for IDH1 R132Q. While this region accommodated the inhibitor in the R132H:NADP(H) complex, these residues would interfere with inhibitor binding to R132Q:NADP(H). Interestingly, unlike in R132Q, these residues didn’t appear to preclude inhibitor binding in WT:NADP(H). Thus, while it is the α10 regulatory segment that precludes inhibitor binding in IDH1 WT, it is instead residues 111-121 that prevent inhibitor binding in R132Q. This suggests that the essentially kinetically identical inhibitory characteristics of IDH1 WT and R132Q ^22^ develop through two very different mechanisms. Importantly, as this loop would not have been readily apparent as a selectivity gate when only examining the IDH1 WT structure, it is only through our R132Q:NADP(H) structure that we were able to identify a possible novel resistance strategy and selectivity handle.

While much effort has been devoted to understanding the unique catalytic and structural features of IDH1 WT versus R132H, our discovery of the unusual kinetic properties of the IDH1 R132Q mutant allowed a valuable opportunity to establish the static and dynamic structural adjustments required to maintain conventional and neomorphic activities within the same active site. Compared to IDH1 R132H, our findings show that the IDH1 R132Q binding pocket and surrounding areas are better primed for substrate binding and hydride transfer steps. Rather than simply acting as a hybrid of WT and R132H, IDH1 R132Q employed unique strategies to yield improved catalytic parameters for both ICT and αKG turnover as compared to R132H. These structural and dynamic discoveries not only highlight mechanistic properties of important tumor drivers, but also identify novel regions that may serve as selectivity handles when designing mutant IDH1 inhibitors requiring increasing selectivity or optimization against resistance mutants.

## Materials and Methods

### Reagent and tools

Dithiothreitol (DTT), isopropyl 1-thio-β-D-galactopyranoside (IPTG), Triton X-100, α-ketoglutaric acid sodium salt (αKG), DL-isocitric acid trisodium salt hydrate, and magnesium chloride (MgCl_2_) were obtained from Fisher Scientific (Hampton, NH). BME was obtained from MP Biomedicals (Santa Ana, CA). β-Nicotinamide adenine dinucleotide phosphate reduced trisodium salt (NADPH), β-Nicotinamide adenine dinucleotide phosphate disodium salt (NADP^+^) and tris(2-carboxyethyl)phosphine) (TCEP) was purchased from Millipore Sigma (Burlington, MA). Nickel-nitrilotriacetic acid (Ni-NTA) resin was obtained from Qiagen (Valencia, CA). Stain free gels (4-12%) were obtained from Bio-Rad Laboratories (Hercules, CA). Protease inhibitor tablets were obtained from Roche Applied Science (Penzberg, Germany). Phenylmethylsulfonyl fluoride (PMSF) salt was purchased from Thermo Scientific (Waltham, MA). The *Escherichia coli* BL21 Gold DE3 strain was used for all protein expression.

### Purification of IDH1 WT and mutant

Human IDH1 WT, R132H, and R132Q homodimers were expressed from a pET-28b(+) plasmid and purified as described previously ^21^ for steady-state kinetic analysis. For pre-steady-state kinetics and HDX-MS experiments, protein was loaded onto a pre-equilibrated (50 mM Tris-HCl 7.5 at 4 °C and 100 mM sodium chloride) Superdex 16/600 size exclusion column (GE Life Sciences, Chicago, IL) following Ni-NTA affinity chromatography to remove any protein aggregate. Protein was eluted with 50 mM Tris-HCl pH 7.5 at 4 °C, 100 mM NaCl, and 1 mM DTT. The fractions were pooled and concentrated for use in pre-steady-state experiments, or pooled and dialyzed in Tris-HCl pH 7.5 at 4 °C, 100 mM NaCl, 20% glycerol, and 1 mM DTT and used immediately for HDX-MS analysis ^31^. For IDH1 R132Q X-ray crystallography experiments, two 1 L cultures of terrific broth supplemented with 50 μg/ml of kanamycin were incubated at 37 °C and 180 rpm until an A_600_ of 0.4 was reached. Cultures were removed and placed onto stir plates and allowed to cool to 25 °C. Expression was induced when cultures reached an A_600_ of 0.8-1.0 with 1 mM IPTG and incubated for an additional 16-18 hours. Cell pellets were harvested and resuspended in lysis buffer (20 mM Tris pH 7.5 at 4 °C, 500 mM NaCl, 0.2% Triton X-100, 5 mM imidazole, 1 mM PMSF, and 5 mM BME). Following cell lysis via sonication, crude lysate was clarified via centrifugation at 14,000 x *g* for one hour. The lysate was loaded on to a pre-equilibrated Ni-NTA column. The column was washed with 100 mL of wash buffer (20 mM Tris pH 7.5 at 4 °C, 500 mM NaCl, 15 mM imidazole, 5 mM BME). Protein was eluted using elution buffer (50 mM Tris pH 7.5 at 4°C, 500 mM NaCl, 500 mM imidazole, 5% glycerol, 10 mM BME). For the NADP(H)-stripped experiments, a buffer containing αKG was passed through the Ni-NTA affinity column prior to elution as described in previous work ^14^. In all cases, eluted protein was loaded onto a HiPrep 26/10 desalting column (GE Healthcare) containing 25 mM Tris pH 7.5 at 20 °C, 500 mM NaCl, 5 mM EDTA, 2 mM DTT, and placed on ice overnight to remove any remaining metals from purification. Fractions containing IDH1 were concentrated (MilliPore Amicon Ultra 15 30 kDa NMWL concentrator) and loaded onto a Superdex 26/600 (GE Healthcare) pre-equilibrated with 20 mM Tris pH 7.5 at 20 °C, 200 mM NaCl, and 2 mM DTT. Fractions containing pure IDH1 were pooled and concentrated to a final concentration of 14-20 mg/mL, flash frozen using liquid nitrogen, and stored at -80 °C. In all cases, the purity of the protein (>95% was confirmed using SDS-PAGE analysis).

### Molecular graphics images

Structure figures were prepared using PyMOL^40^.

### Kinetics assays

To measure steady-state activity of homodimer WT, R132H, and R132Q, only minor modifications were made from previous studies ^21,22^. For the conventional reaction (ICT to αKG), IDH1 buffer (50 mM Tris HCl pH 7.5 at 37 °C, 150 mM NaCl, 10 mM MgCl_2_, 1 mM DTT) and homodimer IDH1 (100 nM IDH1 WT, or 200 nM IDH1 R132H and R132Q), as well as various concentrations of ICT and 200 µM NADP^+^ were preincubated separately for 3 min at 37 °C. Following addition of substrates at 37 °C, the increase of absorbance at 340 nm due to production of NADPH was monitored using an Agilent Cary UV/Vis 3500 spectrophotometer (Santa Clara, CA). For the neomorphic reaction (αKG to D2HG), IDH1 buffer and homodimer mutant IDH1 (200 nM) as well as various concentrations of αKG at pH 7.5 and 200 µM NADPH were separately preincubated for 3 min at 37 °C. Following addition of substrates at 37 °C, the decrease of absorbance at 340 nm due to consumption of NADPH was monitored. The kinetic parameters, which were obtained using at least two unique protein preparations, were determined as described previously ^21,22^.

For the reducing agent inhibition steady-state studies, the conventional reaction conditions described above were repeated except one of three reducing agents (DTT, TCEP, or BME) were added at varying concentrations during the pre-incubation step with the enzyme before substrates were added. Upon obtaining Michaelis-Menten plots at various reducing agent concentrations, the inverse of both *k*_obs_ and substrate concentration were plotted in Lineweaver-Burk analysis.

Single-turnover, pre-steady-state kinetic assays were performed for the neomorphic reaction at 37 ⁰C using an RSM stopped-flow spectrophotometer (OLIS, Atlanta, GA). For the neomorphic reaction, hydride transfer (NADPH to NADP^+^ conversion) was monitored as a change in fluorescence as a function of time via measuring the depletion of NADPH signal by exciting the sample at 340 nm and scanning the emission spectrum from 410 to 460 nm. Final concentrations after mixing were as follows: 40 µM IDH1 R132Q or R132H, 10 µM NADPH, 10 mM αKG (IDH1 R132H) or 0.5 mM αKG (IDH1 R132Q), 50 mM Tris-HCl (pH 7.5 at 37 °C), 150 mM NaCl, 0.1 mM DTT, and 10 mM MgCl_2_. The change in fluorescence as a function of time was fit to a single exponential equation (Y = A_0_e^-kt^) using Graphpad Prism to obtain *k*_obs_. For IDH1 R132H, a higher concentration of αKG (20 mM) was used since 1 mM αKG showed an initial lag.

Single turnover pre-steady-state kinetics were also performed for the conventional reaction at 37 ⁰C to obtain rate constants associated with steps after NADP^+^ binding through hydride transfer using an RSM stopped-flow spectrophotometer. NADPH formation as a function of time was similarly monitored by exciting at 340 nm and scanning the emission spectrum from 410 to 460 nm. Final concentrations after mixing were as follows: 30 µM IDH1 WT or R132Q, 10 µM NADP^+^, 0.5 mM ICT (IDH1 WT) or 1 mM ICT (IDH1 R132Q), 50 mM Tris-HCl (pH 7.5 at 37 °C), 150 mM NaCl, 0.1 mM DTT, and 10 mM MgCl_2_. The change in fluorescence as a function of time was fit to a single exponential equation (Y = A_0_e^-kt^) using Graphpad Prism and *k*_obs_ values were obtained.

Rates associated with NADPH binding corresponding to the first step of the catalytic cycle for the neomorphic reaction were performed as previously described ^14^ using an RSM-stopped flow spectrophotometer (OLIS, Atlanta, Georgia). However, due to low sensitivity of our stopped-flow spectrophotometer, the concentrations of NADPH and IDH1 were increased 10-fold, which in the case of IDH1 WT led to rates too fast to be detected by our instrument (≤100 s^-1^). Therefore, glycerol (40%) and temperature (10 °C) were used to slow NADPH binding rates to IDH1. NADPH binding as a function of time was monitored by exciting at 340 nm and scanning the emission spectrum from 410 to 460 nm. Final concentrations after mixing were as follows: 4 µM IDH1, varying concentration of µM NADP^+^, 100 mM Tris-HCl pH 7.5, 150 mM NaCl, 0.1 mM DTT, 10 mM MgCl_2_, and 40% glycerol. The change in fluorescence as a function of time was fit to a single exponential equation (Y = A_0_e^-kt^) using Graphpad Prism, and *k*_obs_ values were obtained and plotted as a function of NADPH concentration using the equation *k_obs_* = *k_1_*[NADPH] + *k_-1_*. This yielded a linear graph indicating one-step binding, with the slope equal to *k_1_* and the Y-intercept equal to *k_-1_*, though the Y-intercept slope was too high to do so reliably.

Isothermal titration calorimetry (ITC) experiments were conducted at the Sanford Burnham Prebys Protein Production and Analysis Facility using a Low Volume Affinity ITC calorimeter (TA Instruments). For NADPH titrations, experiments were performed at 25 °C in 20 mM Tris pH 7.5, 100 mM NaCl, 10 mM MgCl_2_, and 2 mM BME, injecting 0.25 mM NADPH into the cell containing 0.025 mM for IDH1 WT, 0.025 mM or 0.04 mM IDH1 R132H; and injecting 0.15 mM NADPH into the cell containing 0.034 mM or 0.026 mM IDH1 R132Q. For ICT titrations, experiments were performed at 25 °C in 20 mM Tris pH 7.5, 100 mM NaCl, 10 mM CaCl_2_, and 2 mM BME, injecting 0.6mM ICT into the cell containing 0.12 mM IDH1 R132Q or 0.13 mM IDH1 R132H. Baseline control experiments were performed by injecting the ligand into a cell with buffer only. In all cases, ITC data were analyzed using the Nanoanalyze software package by TA Instruments.

### HDX-MS data collection and analysis

HDX-MS data collection and analysis was performed at the Biomolecular and Proteomics Mass Spectrometry Facility (BPMSF) of the University California San Diego using a Waters Synapt G2Si system with HDX technology (Waters Corporation, Milford, MA) as previously described, again using a sample of IDH1 WT without substrates to be analyzed alongside every experiment to allow experiment to experiment comparisons ^31,41^. Deuterium exchange reactions were conducted using a Leap HDX PAL autosampler (Leap Technologies, Carrboro, NC). The D_2_O buffer was prepared by lyophilizing dialysis buffer (50 mM Tris buffer at pH 7.5 at 4 °C, 100 mM NaCl, and 1 mM DTT) either alone (IDH1:NADP(H) condition) or with the following ligands: for the conventional reaction experiments, IDH1 WT and R132Q were treated with 0.01 mM NADP^+^ and 10 mM ICT (ternary complexes), or with 0.1 mM NADP^+^, 10 mM ICT, and 10 mM CaCl_2_. For the neomorphic reaction experiments, IDH1 R132Q and R132H were treated with 0.1 mM NADPH and 10 mM αKG (ternary complexes), or with 0.1 mM NADPH, 10 mM αKG, and 10 mM CaCl_2_ was also included (quaternary complexes). The buffer was first prepared in ultrapure water and then redissolved in an equivalent volume of 99.96% D_2_O (Cambridge Isotope Laboratories, Inc., Andover, MA) just prior to use. Deuterium exchange measurements were performed in triplicate for every time point (in the order of 0 min, 0.5 min, 1 min, 2 min, 5 min); each run took 30 min to complete. Samples were prepared ∼ 30 min prior to experimental setup and stored at 1 °C until dispensing into reaction vials at the start of the reaction, resulting in samples that were exposed to its deuterium buffer between 2 h (0.5 min timepoint) and 7.5 h (last replicate of the 5 min timepoint) at 1 °C. Protein (4 µL) alone or with ligands were equilibrated for 5 min at the reaction temperature (25 °C) before mixing with D_2_O buffer (56 µL, +/- ligands depending on condition). A solution of 3 M guanidine hydrochloride (50 µL, final pH 2.66) was added to each sample (50 µL) with incubation for 1 min at 1 °C to quench deuterium exchange. The quenched sample (90 µL) was injected into a 100 uL sample loop for in-line digestion at 15 °C using pepsin column (Immobilized Pepsin, Pierce). Peptides were then captured on a BEH C18 Vanguard precolumn and then separated by analytical chromatography (Acquity UPLC BEH C18, 1.7 µm 1.0 × 50 mm, Waters Corporation) over 7.5 min using a 7-85% acetonitrile gradient containing 0.1% formic acid. Samples were then electrosprayed into a Waters Synapt G2Si quadrupole time-of-flight mass spectrometer. Data were collected in the Mobility, ESI+ mode (mass acquisition range = 200–2000 (m/z); scan time = 0.4 s). An infusion of leu-enkephalin (m/z = 556.277) every 30 s (mass accuracy of 1 ppm for calibration standard) was used for continuous lock mass correction.

To identify peptides, data was collected on the mass spectrometer in mobility-enhanced data-independent acquisition (MS^E^), mobility ESI+ mode. Peptide masses were determined from triplicate analyses, and resulting data were analyzed using the ProteinLynx global server (PLGS) version 3.0 (Waters Corporation). We identified peptide masses using a minimum number of 250 ion counts for low energy peptides and 50 ion counts for their fragment ions, with the requirement that peptides had to be larger than 1,500 Da in all cases. Peptide sequence matches were filtered using the following cutoffs: minimum products per amino acid of 0.2, minimum score of 7, maximum MH+ error of 5 ppm, and a retention time RSD of 5%. To ensure high quality, we required that all peptides were present in two of the three experiments. After identifying peptides in PLGS, we then used DynamX 3.0 data analysis software (Waters Corporation) for peptide analysis. Here, relative deuterium uptake for every peptide was calculated via comparison of the centroids of the mass envelopes of the deuterated samples with non-deuterated controls per previously reported methods ^42^, and used to obtain data for coverage maps. Data are represented as mean values +/- SD of the three technical replicates due to processing software limitations, but we note that the LEAP robot provides highly reproducible data for biological replicates. Back-exchange was corrected for in the deuterium uptake values using a global back exchange correction factor (typically ∼25%) determined from the average percent exchange measured in disordered termini of varied proteins ^43^. Significance among differences in HDX data points was assessed using ANOVA analyses and t tests (*p* value cutoff of 0.05) within DECA ^44^. We generated deuterium uptake plots in DECA (github.com/komiveslab/DECA), with data plotted as deuterium uptake (corrected) versus time. An HDX-MS data summary table is shown in Supplementary Table 5, and percent uptake plots are shown in the Source Data files.

### Crystallization

For the NADP(H)-only bound IDH1 R132Q crystals (PDB 8VHC, 8VH9), enzyme (14-20 mg/mL) was incubated on ice with 10 mM NADPH. Crystals of R132Q:NADP(H) were grown via hanging drop vapor diffusion at 4 °C. 2 μL of IDH1 were mixed with 2 μL of well solution containing either 220 mM ammonium sulfate, 100 mM bis-tris pH 6.5, and 20% (w/v) PEG 3350 (PDB ID 8VHC), or well solution containing 200 mM ammonium citrate tribasic pH 7.0 and 26% (w/v) PEG 3350 (8VH9). Though both forms aligned very well and appeared otherwise identical, we feared the citrate buffer could nonetheless promote more substrate-bound-like features due to its structural similarity to isocitrate. Thus, the binary structure crystallized in sulfate was used for all further comparisons and alignments.

IDH1 R132Q crystals containing ICT (8VHD) were grown by first incubating the enzyme at 20 mg/mL with 10 mM NADP^+^, 10 mM CaCl_2,_ and 200 mM DL-isocitric acid at 20°C for 1 h. Then, 2 µL of IDH1 were mixed with 2 μL of well solution containing 100 mM bis-tris propane pH 6.5, 200 mM NaI, and 24% (w/v) PEG 3350 and stored at 4 °C. Crystals were harvested using a nylon-loop and cryo-protected using a solution of 100 mM bis-tris propane pH 6.5, 200 mM NaI, 26%(w/v) PEG 3350, and 20%(v/v) glycerol. Crystals were flash-frozen in liquid nitrogen and stored until data collection.

IDH1 R132Q crystals containing αKG and/or αKG-adducts were generated by incubating enzyme (14-20 mg/mL) on ice with 10 mM NADPH, 20 mM CaCl_2_, 75 mM αKG Fisher Scientific (Hampton, NH) for 1 h. For the 8VHB structure, crystals were grown at 4°C via hanging drop vapor diffusion, where 2 μL of IDH1 were mixed with 2 μL of the well solution containing 200 mM NaSCN and 21%(w/v) PEG 3350. Crystals were cryo-protected using a solution of 20% (v/v) glycerol, 25% (w/v) PEG 3350 and 200 mM NaSCN, and flash-frozen in liquid nitrogen and stored until data collection. For the 8VHA structure, IDH1 R132Q was incubated at 20 °C with 10 mM NADPH, 10 mM CaCl_2_, 10 mM αKG, and then crystals were grown at 4 °C by mixing 2 μL of IDH1 R132Q with 2 μL of well solution containing 160 mM NaNO_3_ and 20% (w/v) PEG 3350. Crystals were harvested using a nylon-loop and cryo-protected in a solution containing 22% (v/v) glycerol and 26% (w/v) PEG 3350.

For IDH1 R132Q crystals containing the NADP-TCEP adduct (8VHE), enzyme (14-20 mg/mL) was incubated on ice with 10 mM NADP^+^, 20mM CaCl_2_, and 75 mM DL-isocitric acid for 1 h. Crystals were grown at 4 °C via hanging drop vapor diffusion, with 1.5 μL of IDH1 mixed with 1.5 μl of well solution containing 200 mM KSCN, 24% (w/v) PEG 6000, and 5 mM TCEP pH 7.4.

### Data collection, processing, and refinement

Data were collected at 100K using synchrotron radiation at the Advanced Photon Source, beamline 24-ID-E or at the Stanford Synchrotron Radiation Lightsource, beamline BL12-2. All datasets were processed with XDS ^45^. Structure solutions were obtained by molecular replacement using PHASER-MR in Phenix ^46,47^. For αKG and/or αKG-adducts (8VHB and 8VHA), isocitrate (8VHD), and NADP-TCEP (8VHE) co-crystals, PDB ensembles of 1T0L ^13^, 4KZO ^14^, and 6PAY ^26^ were used for molecular replacement by generating ensembles using Phenix Ensembler ^46,47^. For IDH1 R132Q apo structures, 1T09 and 4UMX were used as search models. The models were optimized via iterative rounds of refinement in Phenix Refine and manual rebuilding in Coot ^48,49^. Ligand restraints were generated in Phenix eLBOW ^46,47^. Data collection and refinement statistics are summarized in Supplementary Table 6, and a stereo-image of the electron density maps for each new structure are shown in Supplementary Figure 6.

### Calculations

Density functional theory (DFT) calculations ^50^ were carried out to model the NADP-TCEP binding energetics and geometry using the Gaussian 16 suite of programs ^51^. The NADP^+^ was modeled as the nicotinamide ring plus a pendant dihydroxy furan to represent the sugar. The model NADP^+^ and NADP-TCEP adduct were each given a +1 charge. To better model the effects of the solvent, three explicit water molecules were included in calculations on the adducts, distributed at the likeliest sites for hydrogen bonding. The B3LYP ^52^, ωB97XD ^53^, and M06 ^54^ hybrid functionals were used with the cc-pVDZ ^55,56^ and pc-*n* ^57,58^ basis sets, with the latter obtained from the online Basis Set Exchange ^59^. In all of these calculations, implicit solvation was applied using the COSMO model with water as the solvent ^60,61^ and empirical dispersion was added using the D3 version of Grimme’s dispersion along with Becke-Johnson damping ^62,63^. This treatment of solvation effectively models the species as though they were in solution rather than crystalline form. Harmonic frequency analysis was carried out to obtain the vibrational corrections needed to calculate the free energies. Finally, because basis set superposition error can be substantial relative to intermolecular bond energies, the counterpoise correction was applied to our final energies of reaction ^64,65^. The transition state (TS) for the TCEP + NADP binding was identified and confirmed by analysis of the single imaginary vibrational frequency. The DFT calculations for the model NADP-TCEP adduct predicted values of 25° for Δ*θ*_C_ and -11° for Δ*θ*_N_, where the experimental values in the X-ray structure were Δ*θ*_C_ = 29.2° and Δ*θ*_N_ = -1.1° (Supplementary Table 2). For the NADP-αKG adduct, agreement was similar, with DFT predicting Δ*θ*_C_=29° and Δ*θ*_N_= -14° as compared to Δ*θ*_C_=25° and Δ*θ*_N_= -25° in the X-ray structure (Supplementary Table 2). The binding was energetically favored, and appeared to occur without barrier when vibrational effects were included, with a calculated binding energy of 9.4 kcal mol^-1^ at 298 K. However, the calculated free energies indicated that in solution, the entropy decrease would preclude spontaneous binding. Quenching the translational entropy of the species in the crystal may be what allowed the process to occur. We noted that the counterpoise corrections to the transition state and adduct energies were essential, having magnitudes of 7-8 kcal mol^-1^ and comparable to the uncorrected energy differences.

For the dihedral angles, the deviation from planarity Δ*θ* of the NADP pyridine ring in the adduct was reported using the average of two dihedral angles. Numbering the carbon atoms in the ring by convention as shown in Appendix Fig S2, the C-P bond in NADP-TCEP formed at atom 4. The positions of the N atom 1 and the opposite C atom 4 are referenced to the plane defined by the roughly coplanar atoms 2, 3, 5, and 6. The average of the dihedral angles 2-3-5-4 and 6-3-5-4 (Supplementary Fig. 5) was subtracted from 180° to yield Δ*θ*_C_ as a metric for the deviation from planarity of C4, while the average of 3-2-6-1 and 5-2-6-1 subtracted from 180° is used to calculate Δ*θ*_N_ for N1. A sign convention was applied such that if Δ*θ*_C_ and Δ*θ*_N_ had the same sign, the two corners of the ring bend away each other in chair fashion, whereas opposite signs indicate a boat-like conformation. Comparison of the results from the different functionals and basis sets showed very little difference in the geometry. Optimized geometries obtained with the pc-2 basis set on a smaller geometry (omitting sugar and explicit waters) were not significantly different from those obtained with pc-1, so we chose to report the B3LYP/pc-1 results here, with the sugar and explicit waters included (Supplementary Table 2). An additional geometry optimization was run on the NADP-αKG adduct with two explicit waters and a -2 charge, employing the aug-pc-1 basis set ^57,66^ to obtain the diffuse functions necessary to adequately model anions.

## Data availability

Crystallographic data and protein structure coordinates have been deposited with the Protein Data Bank (PDB) public repository. Output files from the computational work are available at the ioChem-BD database (https://doi.org/10.19061/iochem-bd-6-320). HDX-MS data will be uploaded to a MASSIVE repository with accession number prior to manuscript publication. Extended Data Figs. 1-10 and Supplementary information are provided as separate documents. All deuterium uptake plots are provided in the Source Data files.

## Acknowledgements

This work was funded by a Research Scholar Grant, RSG-19-075-01-TBE, from the American Cancer Society (C.D.S.), National Institutes of Health R35 GM137773 (C.D.S.), MARC 1 T34 GM149430 (C.D.S.), MARC 5T34GM008303 (SDSU), and IMSD 5R25GM058906 (SDSU), as well as the California Metabolic Research Foundation (SDSU) and the Rees-Steely Research Foundation (E.A.). The HDX-MS core of the UCSD BPMSF is supported by NIH shared instrumentation grant S10 OD0016234. The Sanford Burnham Prebys Protein Production and Analysis Facility is supported by NCI Cancer Center Support Grant P30 CA030199. The Northeastern Collaborative Access Team beamlines are funded by NIH/NIGMS (P30GM124165) and the Eiger 16M detector at the 24-ID-E beam line is funded by a NIH-ORIP HEI grant (S10OD021527). The Advanced Photon Source is a U.S. Department of Energy (DOE) Office of Science User Facility operated for the DOE Office of Science by Argonne National Laboratory under Contract No. DE-AC02-06CH11357. Use of the Stanford Synchrotron Radiation Lightsource, SLAC National Accelerator Laboratory, is supported by the U.S. DOE Office of Science, Office of Basic Energy Sciences under Contract No. DE-AC02-76SF00515. The SSRL Structural Molecular Biology Program is supported by the DOE Office of Biological and Environmental Research, and by NIH/NIGMS (P30GM133894). The content is solely the responsibility of the authors and does not necessarily represent the official views of the National Institutes of Health.

## Author contributions

Matthew Mealka: investigation, data curation, software, formal analysis, validation, methodology, writing – review and editing; Nicole Sierra: investigation, data curation, software, formal analysis, visualization, methodology, writing – review and editing; Diego Avellaneda Matteo: investigation, data curation, software, formal analysis, validation, methodology, visualization writing – review and editing; Elene Albekioni: investigation, data curation, software, formal analysis, validation, methodology, visualization, writing – review and editing; Rachel Khoury: investigation, data curation, software, formal analysis, visualization, methodology, writing – review and editing; Timothy Mai: investigation, software, formal analysis, validation, visualization, writing – review and editing; Brittany Conley: investigation, software, formal analysis, validation, visualization, writing – review and editing; Nalani J. Coleman: investigation, data curation, software, formal analysis, visualization, methodology, writing – review and editing; Kaitlyn A. Sabo: investigation, data curation, software, formal analysis, visualization, methodology, writing – review and editing; Elizabeth A. Komives: funding acquisition, writing – review and editing; Andrey A. Bobkov: data curation, software, formal analysis, validation, methodology, writing – review and editing; Andrew L. Cooksy: investigation, data curation, software, formal analysis, validation, visualization, methodology, writing – review and editing; Steve Silletti: data curation, methodology, formal analysis, validation, writing – review and editing; Jamie Schiffer: investigation, data curation, visualization, software, formal analysis, methodology, writing – review and editing; Tom Huxford: conceptualization, formal analysis, supervision, visualization, writing – review and editing; Christal D. Sohl: conceptualization, formal analysis, data curation, supervision, visualization, funding acquisition, writing – original draft, project administration, writing – review and editing.

## References

1. Dang, L. et al. Cancer-associated IDH1 mutations produce 2-hydroxyglutarate. Nature 462, 739–744 (2009).

2. Leonardi, R., Subramanian, C., Jackowski, S. & Rock, C. O. Cancer-associated isocitrate dehydrogenase mutations inactivate NADPH-dependent reductive carboxylation. J Biol Chem 287, 14615–20 (2012).

3. Pietrak, B. et al. A tale of two subunits: how the neomorphic R132H IDH1 mutation enhances production of alphaHG. Biochemistry 50, 4804–12 (2011).

4. Chowdhury, R. et al. The oncometabolite 2-hydroxyglutarate inhibits histone lysine demethylases. EMBO Rep 12, 463–9 (2011).

5. Figueroa, M. E. et al. Leukemic IDH1 and IDH2 mutations result in a hypermethylation phenotype, disrupt TET2 function, and impair hematopoietic differentiation. Cancer Cell 18, 553–67 (2010).

6. Dang, L., Yen, K. & Attar, E. C. IDH mutations in cancer and progress toward development of targeted therapeutics. Ann Oncol 27, 599–608 (2016).

7. Cleven, A. H. G. et al. IDH1 or -2 mutations do not predict outcome and do not cause loss of 5-hydroxymethylcytosine or altered histone modifications in central chondrosarcomas. Clin Sarcoma Res 7, 8 (2017).

8. Cerami, E. et al. The cBio cancer genomics portal: an open platform for exploring multidimensional cancer genomics data. Cancer Discov 2, 401–4 (2012).

9. Gao, J. et al. Integrative analysis of complex cancer genomics and clinical profiles using the cBioPortal. Sci Signal 6, pl1 (2013).

10. Adeva, J. Current development and future perspective of IDH1 inhibitors in cholangiocarcinoma. Liver Canc Intl 3, 17–31 (2022).

11. Tangella, A. V., Gajre, A. & Kantheti, V. V. Isocitrate dehydrogenase 1 mutation and ivosidenib in patients with acute myeloid leukemia: a comprehensive review. Cureus 15, e44802 (2023).

12. Sharma, N. et al. Isocitrate dehydrogenase mutations in gliomas: A review of current understanding and trials. Neurooncol Adv 5, vdad053 (2023).

13. Xu, X. et al. Structures of human cytosolic NADP-dependent isocitrate dehydrogenase reveal a novel self-regulatory mechanism of activity. J Biol Chem 279, 33946–57 (2004).

14. Rendina, A. R. et al. Mutant IDH1 enhances the production of 2-hydroxyglutarate due to its kinetic mechanism. Biochemistry 52, 4563–77 (2013).

15. Bleeker, F. E. et al. IDH1 mutations at residue p.R132 (IDH1(R132)) occur frequently in high-grade gliomas but not in other solid tumors. Hum Mutat 30, 7–11 (2009).

16. Balss, J. et al. Analysis of the IDH1 codon 132 mutation in brain tumors. Acta Neuropathol 116, 597–602 (2008).

17. Borger, D. R. et al. Frequent mutation of isocitrate dehydrogenase (IDH)1 and IDH2 in cholangiocarcinoma identified through broad-based tumor genotyping. Oncologist 17, 72–9 (2012).

18. Mardis, E. R. et al. Recurring mutations found by sequencing an acute myeloid leukemia genome. N Engl J Med 361, 1058–66 (2009).

19. Yan, H. et al. IDH1 and IDH2 mutations in gliomas. N Engl J Med 360, 765–73 (2009).

20. Pusch, S. et al. D-2-Hydroxyglutarate producing neo-enzymatic activity inversely correlates with frequency of the type of isocitrate dehydrogenase 1 mutations found in glioma. Acta Neuropathol Commun 2, 19 (2014).

21. Avellaneda Matteo, D., et al. Molecular mechanisms of isocitrate dehydrogenase 1 (IDH1) mutations identified in tumors: The role of size and hydrophobicity at residue 132 on catalytic efficiency. J Biol Chem 292, 7971–7983 (2017).

22. Avellaneda Matteo, D., et al. Inhibitor potency varies widely among tumor-relevant human isocitrate dehydrogenase 1 mutants. Biochem J 475, 3221–3238 (2018).

23. Hirata, M. et al. Mutant IDH is sufficient to initiate enchondromatosis in mice. Proc Natl Acad Sci U S A 112, 2829–34 (2015).

24. Casadevall, G., Duran, C. & Osuna, S. AlphaFold2 and Deep Learning for Elucidating Enzyme Conformational Flexibility and Its Application for Design. JACS Au 3, 1554–1562 (2023).

25. Seery, V. L. & Farrell, H. M. J. Spectroscopic evidence for ligand-induced conformational change in NADP+:isocitrate dehydrogenase. J Biol Chem 265, 17644–17648 (1990).

26. Roman, J. V., Melkonian, T. R., Silvaggi, N. R. & Moran, G. R. Transient-state analysis of human isocitrate dehydrogenase I: accounting for the interconversion of active and non-active conformational states. Biochemistry 58, 5366–5380 (2019).

27. Herold, R. A., Reinbold, R., Schofield, C. J. & Armstrong, F. A. NADP(H)-dependent biocatalysis without adding NADP(H). Proc Natl Acad Sci U S A 120, e2214123120 (2023).

28. Farrell, H. M., Jr., Deeney, J. T., Hild, E. K. & Kumosinski, T. F. Stopped flow and steady state kinetic studies of the effects of metabolites on the soluble form of NADP+:isocitrate dehydrogenase. J Biol Chem 265, 17637–43 (1990).

29. Yang, B., Zhong, C., Peng, Y., Lai, Z. & Ding, J. Molecular mechanisms of ‘off-on switch’ of activities of human IDH1 by tumor-associated mutation R132H. Cell Res 20, 1188–200 (2010).

30. Chambers, J. M. et al. Water networks and correlated motions in mutant isocitrate dehydrogenase 1 (IDH1) are critical for allosteric inhibitor binding and activity. Biochemistry 59, 479–490 (2020).

31. Sabo, K. A. et al. Capturing the dynamic conformational changes of human isocitrate dehydrogenase 1 (IDH1) upon ligand and metal binding using hydrogen-deuterium exchange mass spectrometry. Biochemistry 62, 1145–1159 (2023).

32. Sanli, G., Dudley, J. I. & Blaber, M. Structural biology of the aldo-keto reductase family of enzymes: catalysis and cofactor binding. Cell Biochem Biophys 38, 79–101 (2003).

33. Paidimuddala, B., Mohapatra, S. B., Gummadi, S. N. & Manoj, N. Crystal structure of yeast xylose reductase in complex with a novel NADP-DTT adduct provides insights into substrate recognition and catalysis. FEBS J 285, 4445–4464 (2018).

34. Hammes-Schiffer, S. Hydrogen tunneling and protein motion in enzyme reactions. Acc Chem Res 39, 93–100 (2006).

35. Meijers, R. & Cedergren-Zeppezauer, E. A variety of electrostatic interactions and adducts can activate NAD(P) cofactors for hydride transfer. Chem Biol Interact 178, 24–28 (2009).

36. Plapp, B. V. & Ramaswamy, S. Atomic-resolution structures of horse liver alcohol dehydrogenase with NAD(+) and fluoroalcohols define strained Michaelis complexes. Biochemistry 51, 4035–4048 (2012).

37. Patel, S. M. et al. Cautionary tale of using tris(alkyl)phosphine reducing agents with NAD+-dependent enzymes. Biochemistry 59, 3285–3289 (2020).

38. Xie, X. et al. Allosteric mutant IDH1 inhibitors reveal mechanisms for IDH1 mutant and isoform selectivity. Structure 25, 506–513 (2017).

39. Lin, J. et al. Discovery and optimization of quinolinone derivatives as potent, selective, and orally bioavailable mutant isocitrate dehydrogenase 1 (mIDH1) inhibitors. J Med Chem 62, 6575–6596 (2019).

40. Schrödinger, LLC. The PyMOL Molecular Graphics System, Version 2.5.2.

41. Peacock, R. B., Davis, J. R., Markwick, P. R. L. & Komives, E. A. Dynamic consequences of mutation of tryptophan 215 in thrombin. Biochemistry 57, 2694–2703 (2018).

42. Wales, T. E., Fadgen, K. E., Gerhardt, G. C. & Engen, J. R. High-speed and high-resolution UPLC separation at zero degrees Celsius. Anal Chem 80, 6815–6820 (2008).

43. Ramsey, K. M., Dembinski, H. E., Chen, W., Ricci, C. G. & Komives, E. A. DNA and IκBα both induce long-range conformational changes in NFκB. J Mol Biol 429, 999–1008 (2017).

44. Lumpkin, R. J. & Komives, E. A. DECA, a comprehensive, automatic post-processing program for HDX-MS data. Mol Cell Proteomics 18, 2516–2523 (2019).

45. Kabsch, W. XDS. Acta Crystallogr D Biol Crystallogr 66, 125–132 (2010).

46. Liebschner, D. et al. Macromolecular structure determination using X-rays, neutrons and electrons: recent developments in ıt Phenix. Acta Crystallographica Section D 75, 861–877 (2019).

47. Adams, P. D. et al. PHENIX: a comprehensive Python-based system for macromolecular structure solution. Acta Crystallogr D Biol Crystallogr 66, 213–21 (2010).

48. Emsley, P., Lohkamp, B., Scott, W. G. & Cowtan, K. Features and development of Coot. Acta Crystallogr D Biol Crystallogr 66, 486–501 (2010).

49. Emsley, P. & Cowtan, K. Coot: model-building tools for molecular graphics. Acta Crystallogr D Biol Crystallogr 60, 2126–32 (2004).

50. Hohenberg, P. & Kohn, W. Inhomogeneous Electron Gas. Phys. Rev. 136, B864–B871 (1964).

51. Frisch, M. J. et al. Gaussian 16 Rev. C.01. (2016).

52. Becke, A. D. Density-functional thermochemistry. III. The role of exact exchange. J Chem Phys 98, 5648–5652 (1993).

53. Chai, J.-D. & Head-Gordon, M. Long-range corrected hybrid density functionals with damped atom–atom dispersion corrections. Phys. Chem. Chem. Phys. 10, 6615–6620 (2008).

54. Zhao, Y. & Truhlar, D. G. The M06 suite of density functionals for main group thermochemistry, thermochemical kinetics, noncovalent interactions, excited states, and transition elements: two new functionals and systematic testing of four M06-class functionals and 12 other functionals. Theor Chem Acc 120, 215–241 (2008).

55. Dunning, T. H., Jr. Gaussian basis sets for use in correlated molecular calculations. I. The atoms boron through neon and hydrogen. J Chem Phys 90, 1007–1023 (1989).

56. Wilson, A. K., van Mourik, T. & Dunning, T. H. Gaussian basis sets for use in correlated molecular calculations. VI. Sextuple zeta correlation consistent basis sets for boron through neon. J Mol Struct 388, 339–349 (1996).

57. Jensen, F. Polarization consistent basis sets: Principles. J Chem Phys 115, 9113–9125 (2001).

58. Jensen, F. & Helgaker, T. Polarization consistent basis sets. V. The elements Si-Cl. J Chem Phys 121, 3463–70 (2004).

59. Pritchard, B. P., Altarawy, D., Didier, B., Gibson, T. D. & Windus, T. L. New Basis Set Exchange: An Open, Up-to-Date Resource for the Molecular Sciences Community. J. Chem. Inf. Model. 59, 4814–4820 (2019).

60. Barone, V. & Cossi, M. Quantum calculation of molecular energies and energy gradients in solution by a conductor solvent model. J. Phys. Chem. A 102, 1995–2001 (1998).

61. Cossi, M., Rega, N., Scalmani, G. & Barone, V. Energies, structures, and electronic properties of molecules in solution with the C-PCM solvation model. J Comput Chem 24, 669–681 (2003).

62. Grimme, S., Antony, J., Ehrlich, S. & Krieg, H. A consistent and accurate ab initio parametrization of density functional dispersion correction (DFT-D) for the 94 elements H-Pu. J Chem Phys 132, 154104 (2010).

63. Grimme, S., Ehrlich, S. & Goerigk, L. Effect of the damping function in dispersion corrected density functional theory. J Comput Chem 32, 1456–1465 (2011).

64. Boys, S. F. & Bernardi, F. The calculation of small molecular interactions by the differences of separate total energies. Some procedures with reduced errors. Mol Phys 19, 553–566 (1970).

65. Simon, S., Duran, M. & Dannenberg, J. J. How does basis set superposition error change the potential surfaces for hydrogen-bonded dimers? J Chem Phys 105, 11024–11031 (1996).

66. Jensen, F. Polarization consistent basis sets. III. The importance of diffuse functions. J Chem Phys 117, 9234–9240 (2002).

