## Extended Data for "Active site remodeling in tumor-relevant IDH1 mutants drives distinct kinetic features and potential resistance mechanisms"

#### Extended Data Table of Contents:

Extended Data Figs. 1-10

Extended Data References

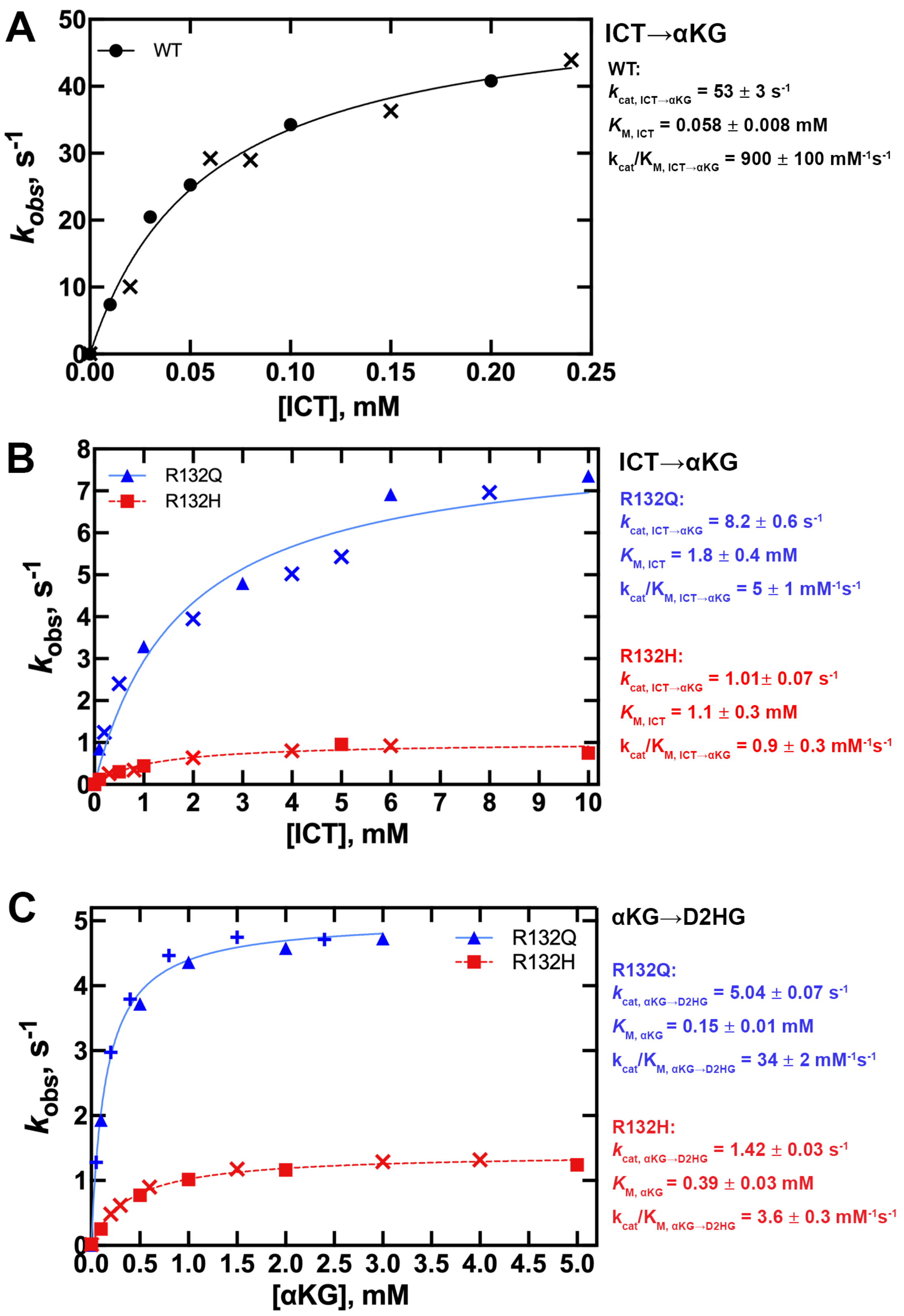


**Extended Data Fig. 1. Steady-state kinetics analyses for IDH1 WT, R132Q, and R132H homodimers.** Steady-state kinetic parameters of the conventional and neomorphic reactions were measured as a function of varying substrate concentration. At least two protein preparations, indicated via different symbols, were used to measure the observed rate constants ($k_{obs}$), which were determined from the linear portion of plots of substrate concentration versus time. Each point in the curve represents a single replicate. A) The conventional reaction catalyzed by IDH1 WT. B) The conventional reaction catalyzed by IDH1 R132Q (blue) and IDH1 R132H (red). C) The neomorphic reaction catalyzed by IDH1 R132Q (blue) and IDH1 R132H (red).

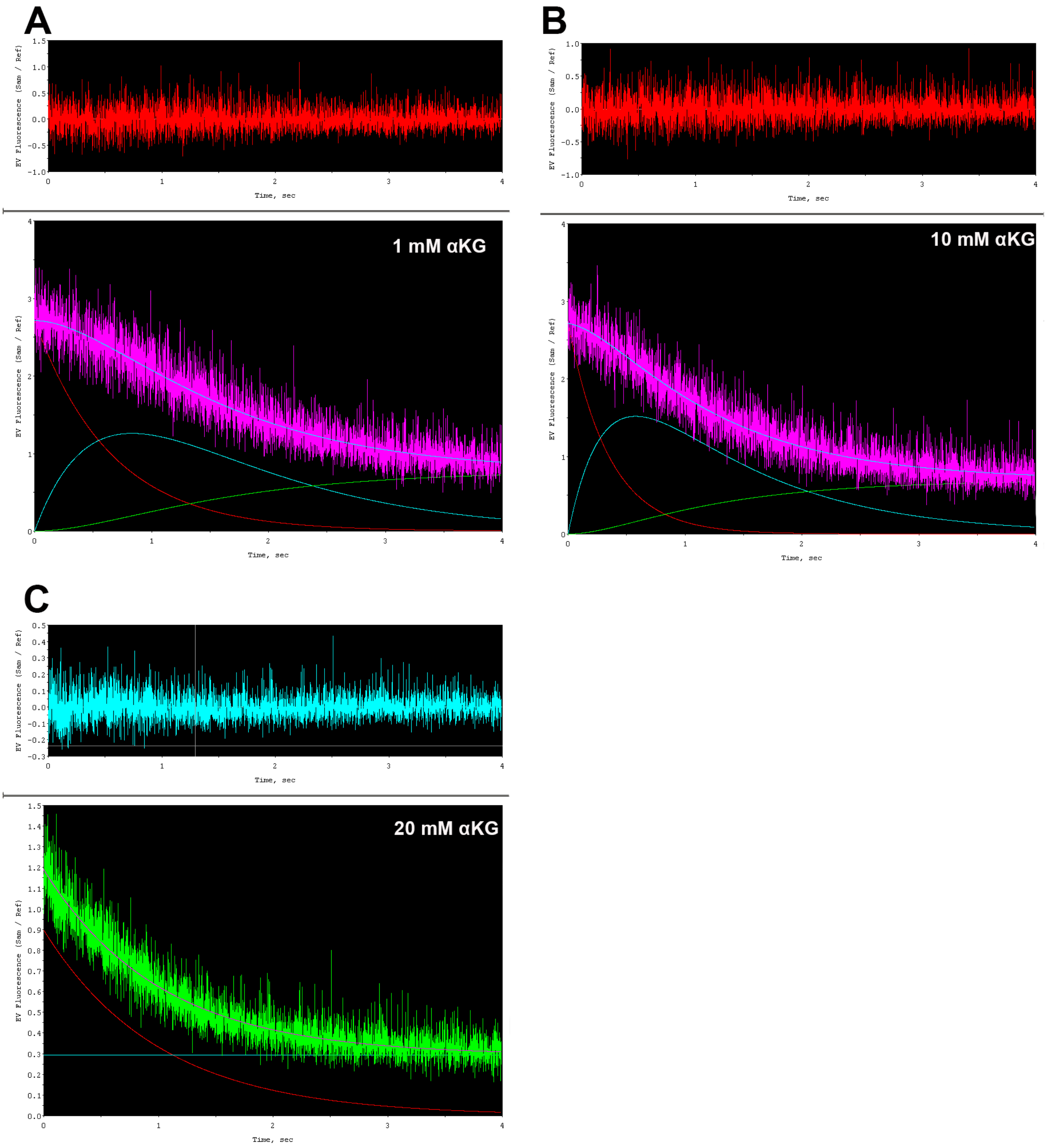


**Extended Data Fig. 2. Lag formation when monitoring hydride transfer by IDH1 R132H upon catalyzing the neomorphic reaction.** A) A notable lag in NADPH consumption during IDH1 R132H catalysis is seen upon treatment of 1 mM αKG. B) The lag was lessened upon increasing substrate concentration to 10 mM αKG. C) The lag was finally eliminated upon using 20 mM αKG. Data in (A) and (B) were fit to a double exponential equation and data in (C) were fit to a single exponential equation, with the residuals shown (top) to assess goodness of fit.

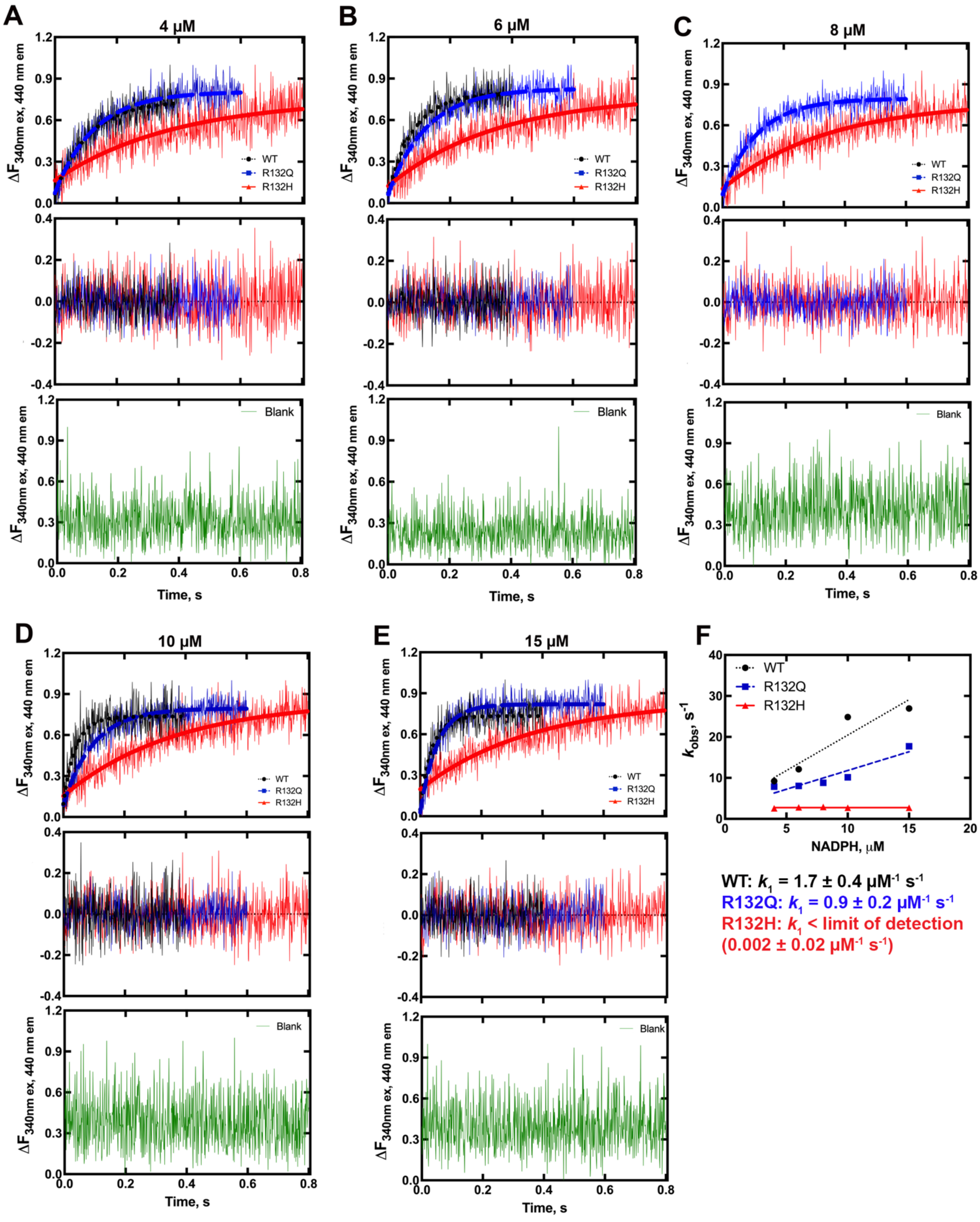


**Extended Data Fig. 3. Rates of NADPH binding to IDH1 WT, R132Q, and R132H.** A-E) NADPH binding was monitored at 10 ºC and 40% glycerol to slow the binding reaction. The change in fluorescence was fit to a single exponential equation (top plot) and residuals (middle plot) were obtained to assess goodness of fit. A control experiment lacking enzyme is shown in the bottom plot (in green). A) 4 µM NADPH. B) 6 µM NADPH. C) 8 µM NADPH. D) 10 µM NADPH. E) 15 µM NADPH. E) The $k_{obs}$ values were plotted as a function of NADPH concentration to yield a linear progression, indicating one-step binding for NADPH.

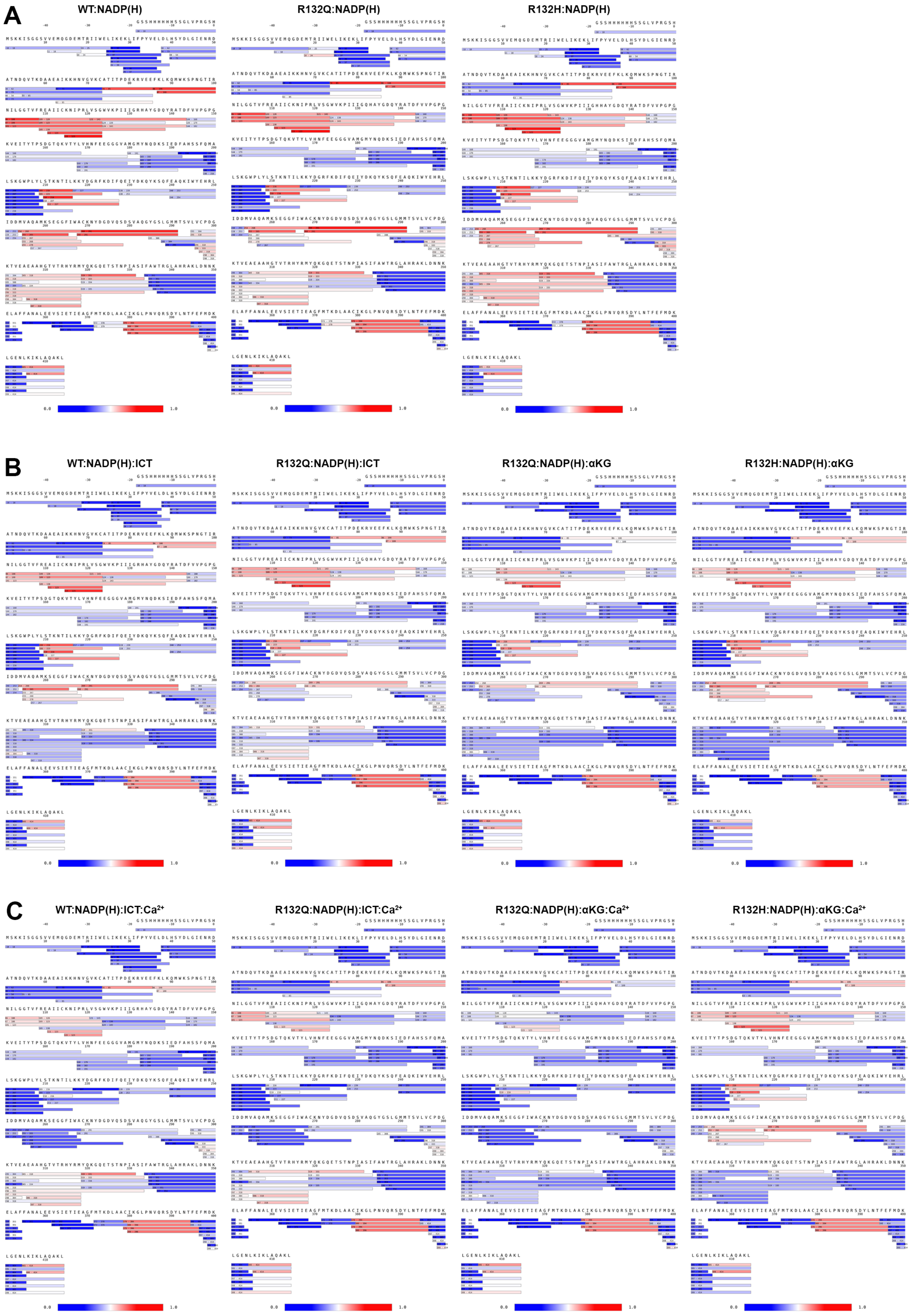
A
WT:NADP(H)
R132Q:NADP(H)
R132H:NADP(H)
B
WT:NADP(H):ICT
R132Q:NADP(H):ICT
R132Q:NADP(H):αKG
R132H:NADP(H):αKG
C
WT:NADP(H):ICT:Ca2+
R132Q:NADP(H):ICT:Ca2+
R132Q:NADP(H):αKG:Ca2+
R132H:NADP(H):αKG:Ca2+
0.0
1.0

**Extended Data Fig. 4. Coverage maps for IDH1 WT, R132Q, and R132H under three unique conditions.** Deuterium uptake is shown as a gradient between highest uptake (red) to lowest uptake (blue). A) The WT, R132H, and R132Q binary form served as a baseline comparison. B) IDH1 WT and R132Q were treated with $NADP^+$ and ICT, and IDH1 R132Q and R132H were treated with NADPH and αKG (ternary complexes). C) WT and mutant IDH1 were treated as in B), except $CaCl_2$ was also included (quaternary complexes).

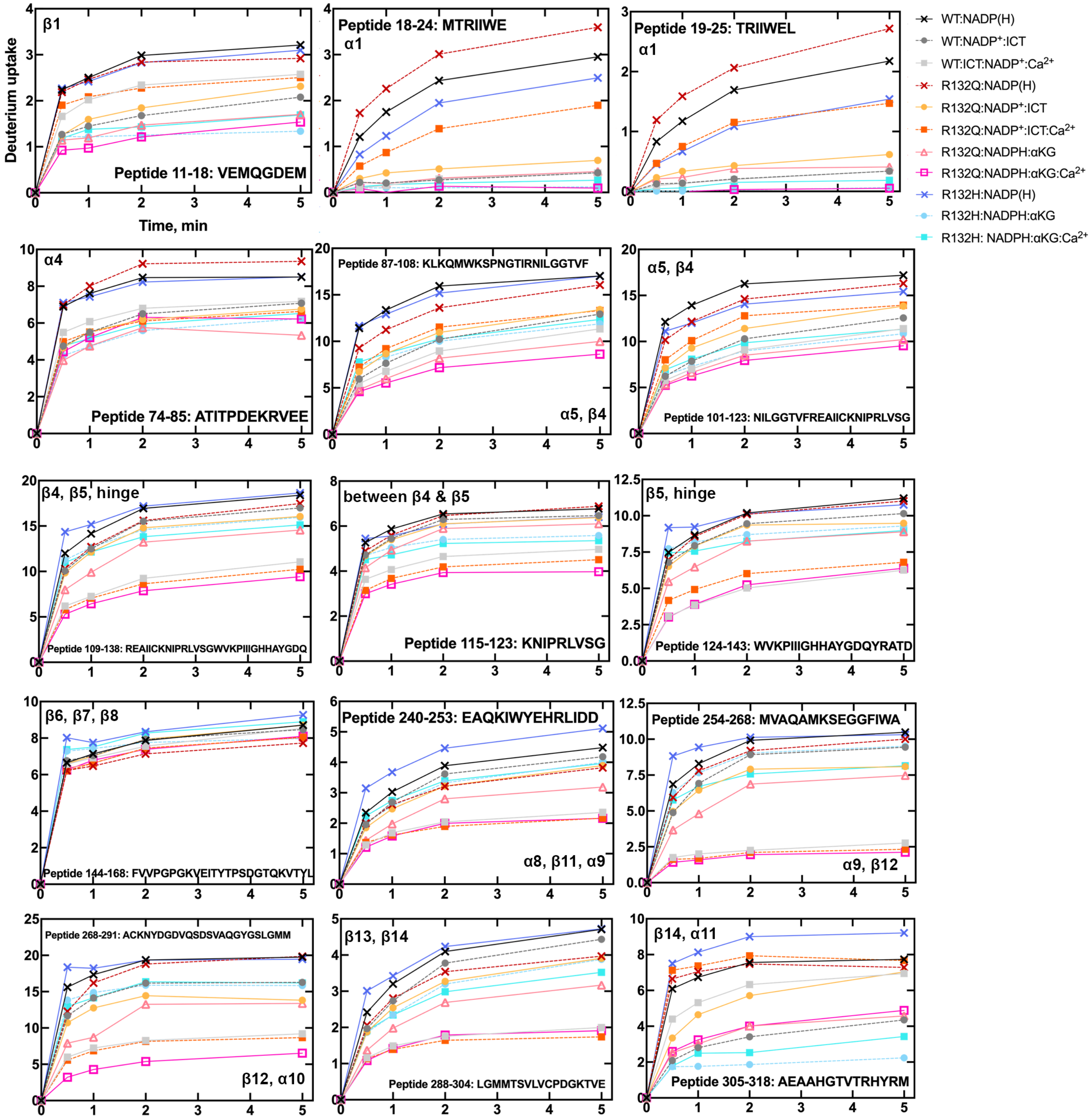


**Extended Data Fig. 5. Deuterium uptake plots.** Back-exchange corrected deuterium uptake was plotted against time for peptides of interest. Conditions are indicated in the legend. In the IDH1:NADP(H) conditions, any cofactors that bound during expression and purification were not removed, though no cofactor incubation step was added. Secondary structures associated with the peptides are indicated.

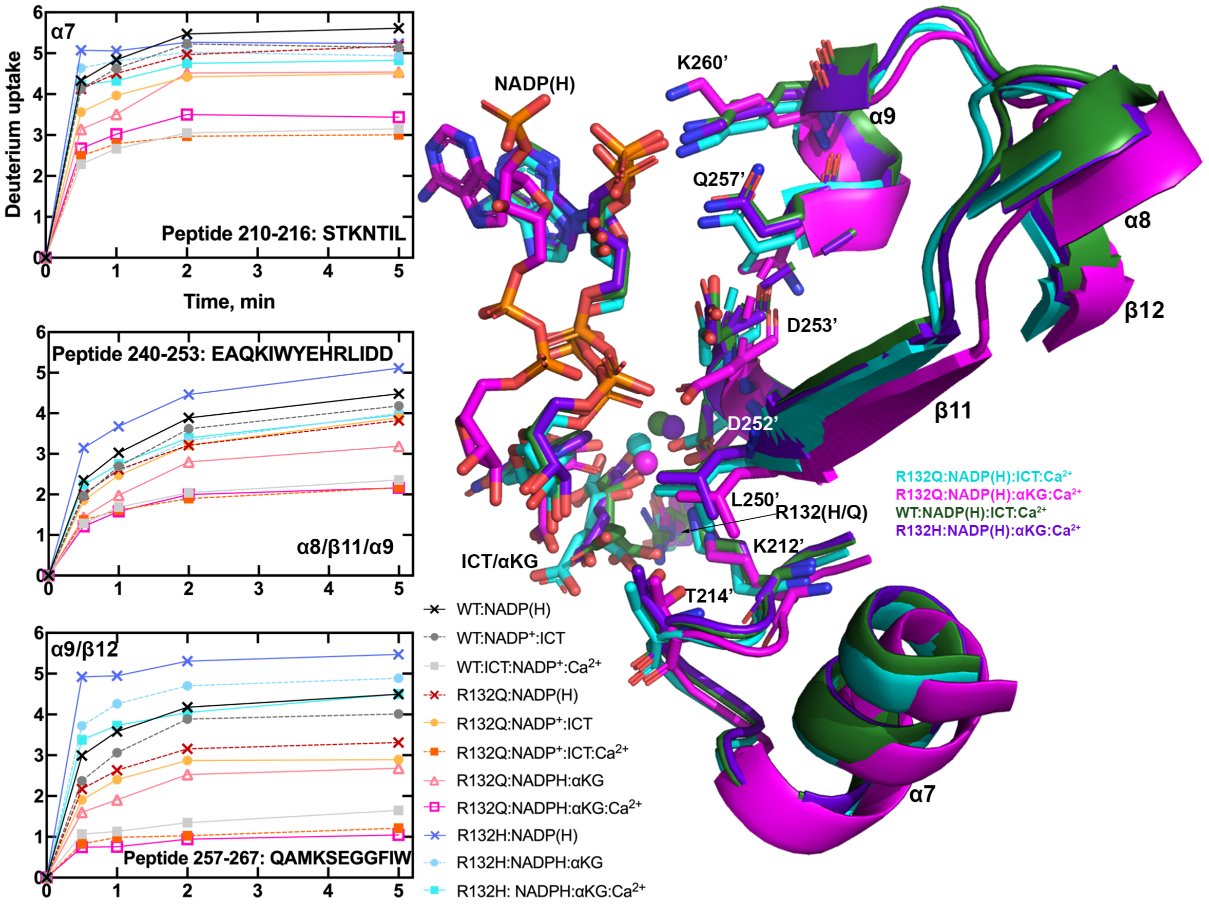


**Extended Data Fig. 6. Peptides near the active site reach deuterium uptake equilibrium faster for IDH1 R132Q and WT relative to R132H upon binding substrates.** Deuterium uptake plots for peptides containing chain B residues within 4 Å of the bound NADP(H) and ICT/αKG molecules (residues K212', T214', L250', D252', D253', Q257', and K260') are shown on the left. Secondary structure associated with these three peptides are shown on the right for ICT-bound R132Q and WT [1], and αKG-bound R132Q and R132H [2]. Secondary structures associated with the peptides are indicated.

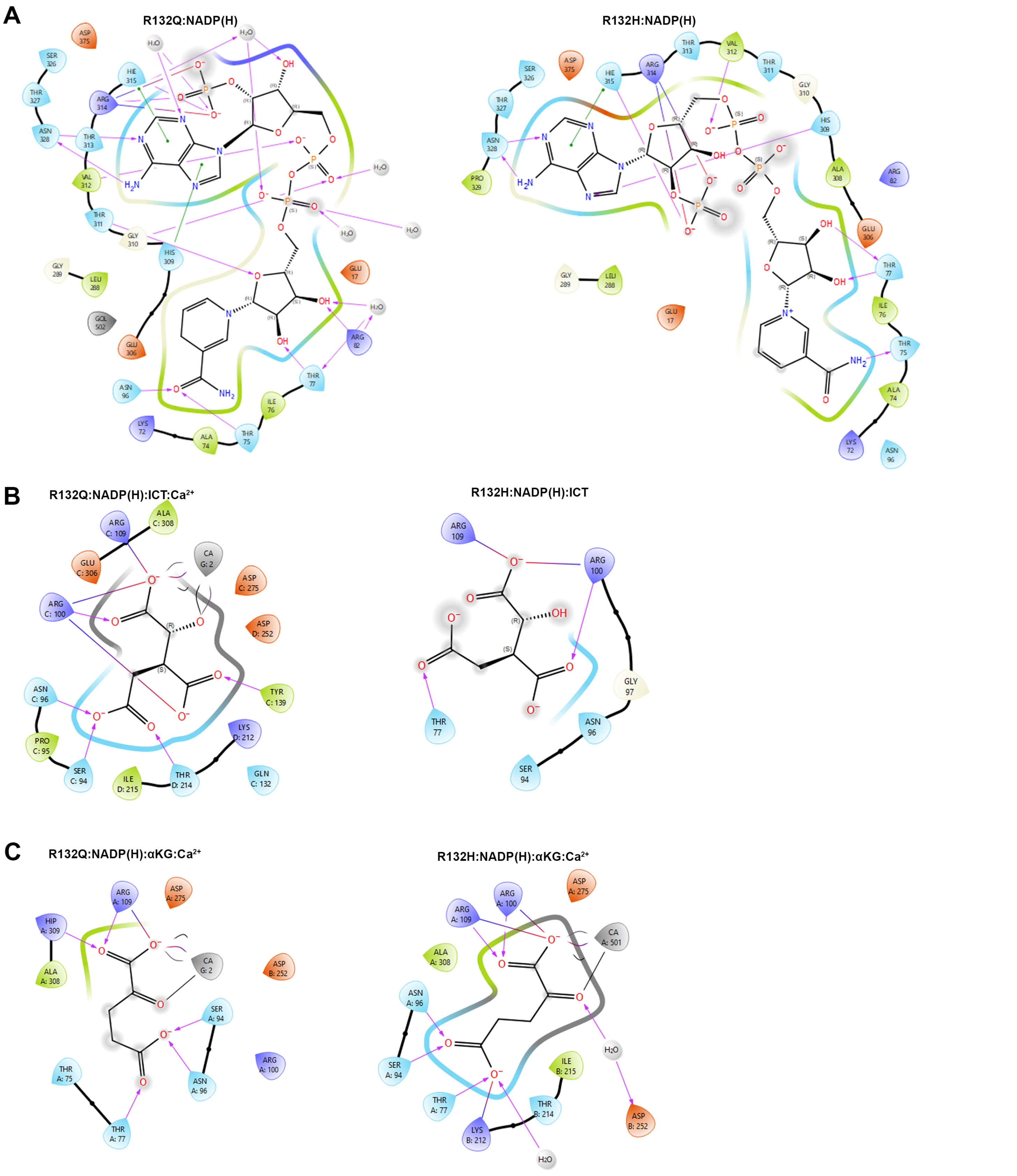


**Extended Data Fig. 7. Ligand interaction diagrams highlighting hydrogen bonding interactions.** In all cases, the residues are shown as guitar picks, with the pointed end signifying the side chain and the rounded end representing the backbone. A) Ligand interaction diagram between NADP(H) and IDH1 R132Q and waters in the cofactor binding pocket for the binary R132Q:NADP(H) structure (from Fig 5A). Hydrogen bonding interactions are shown with pink lines while pi stacking interactions are shown with green. Residues within 4 Å of NADP(H) are shown. In comparison, corresponding hydrogen bonding interactions of NADP(H) are also shown in the previously solved R132H:NADP(H) structure [3]. B) Ligand interaction diagram between ICT and IDH1 R132Q and $Ca^{2+}$ in the R132Q binding pocket for the R132Q:NADP(H):ICT:$Ca^{2+}$ structure. Hydrogen bonding interactions are shown with pink arrows while charge interactions are shown with red-blue lines. Residues within 4 Å of ICT are shown. In comparison, corresponding hydrogen bonding interactions of NADP(H) are also shown in R132H:NADP(H):ICT. C) Ligand interaction diagram between αKG and IDH1 R132Q and $Ca^{2+}$ in the R132Q binding pocket for the R132Q:NADP(H):αKG:$Ca^{2+}$ structure (PDB 8VHB). Hydrogen bonding interactions are shown with pink arrows while charge interactions are indicated with red-blue lines. Residues within 4 Å of αKG are shown. In comparison, corresponding hydrogen bonding interactions of NADP(H) are also shown in the previously solved R132H:NADP(H):αKG:$Ca^{2+}$ structure [2].

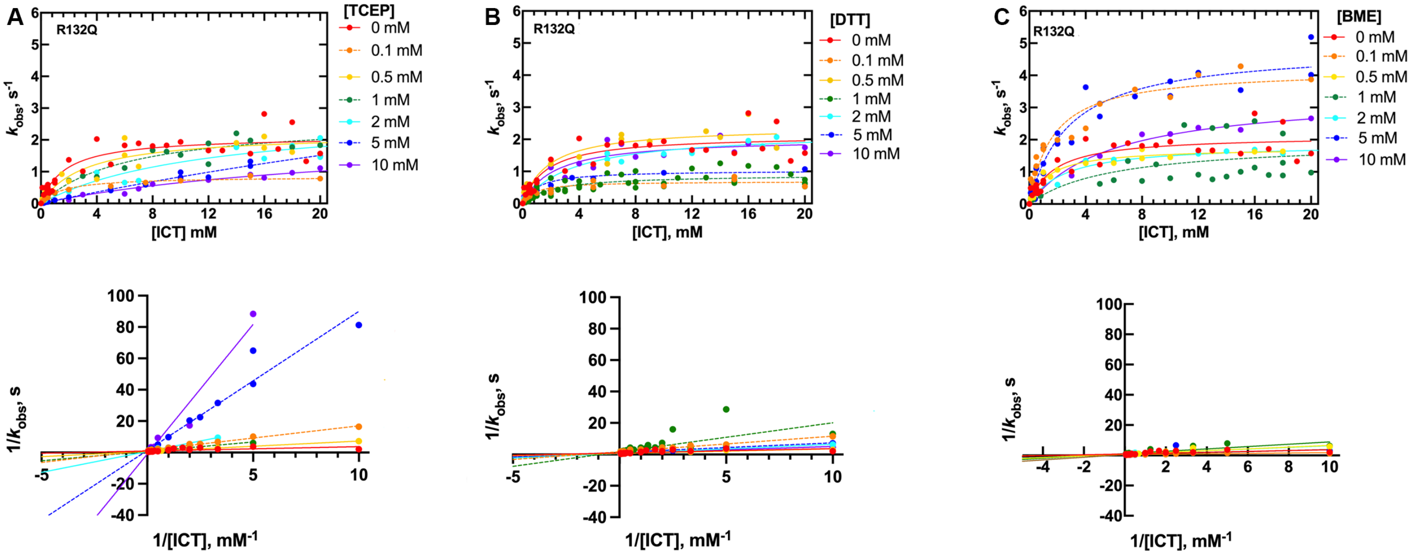


**Extended Data Fig. 8. TCEP treatment competitively inhibits the forward reaction catalyzed by IDH1 R132Q.** Steady-state kinetic parameters of the conventional reaction by IDH1 R132Q were measured as a function of varying ICT concentration upon challenge with one of three reducing agents. Top: At least two protein preparations were used to measure the observed rate constants ($k_{obs}$), which were determined from the linear portion of plots of substrate concentration versus time. Bottom: Lineweaver-Burk analysis was performed by plotting $1/k_{obs}$ *vs* 1/[ICT]. In both plots, each point represents a single replicate. A) IDH1 R132Q catalysis upon increasing concentrations with TCEP. Only treatment with TCEP showed notable, dose-dependent inhibition, with features of competitive inhibition. B) IDH1 R132Q catalysis upon increasing concentrations with DTT. C) IDH1 R132Q catalysis upon increasing concentrations with BME. A table of the kinetic parameters are shown in Supplementary Table 4.

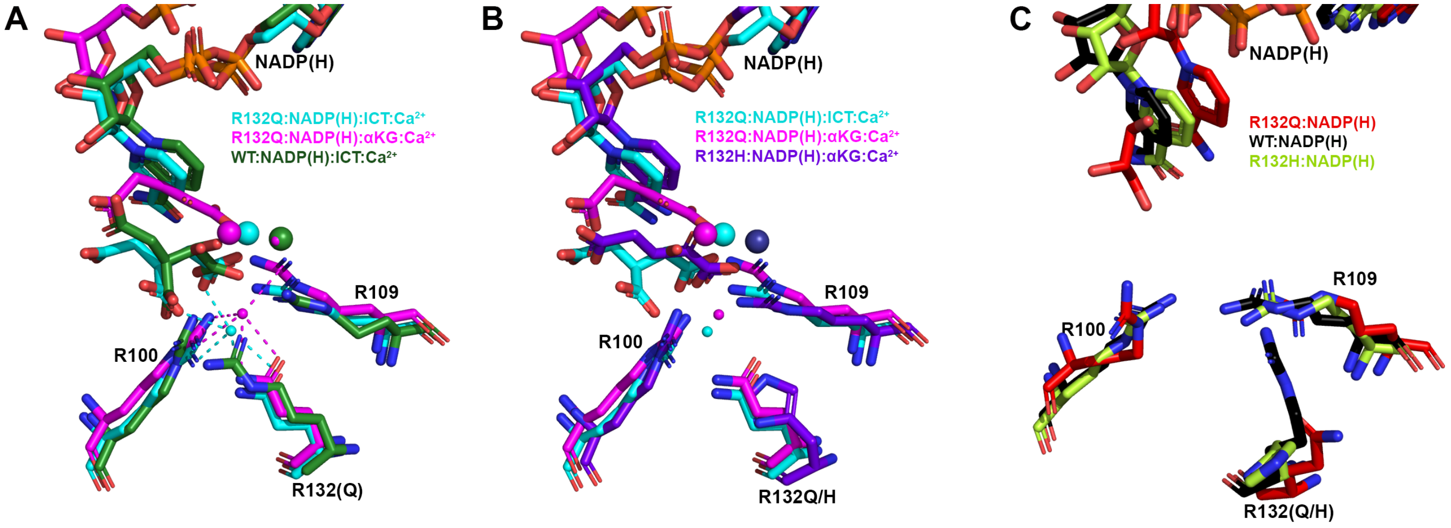


**Extended Data Fig. 9. IDH1 R132Q employs an active site water molecule to mimic polar interactions normally provided by residue R132.** Dimer-based alignments of the active site are shown that highlight residues R100, R109, and R132(Q/H), which play an important role in hydrogen binding to ICT in the absence of mutation at residue R132. A) Both ICT-bound and αKG-bound R132Q structures featured an active site water (small spheres) that helped coordinate ICT, though this water molecule was absent in the quaternary WT structure [1]. A second water molecule localized to the position of the calcium ion (large spheres) in the IDH1 WT structure. B) The quaternary R132H structure [2] did not contain a coordinating active site water. C) The binary NADP(H) bound forms of R132Q, R132H [3], and WT [1] lacked this active site water.

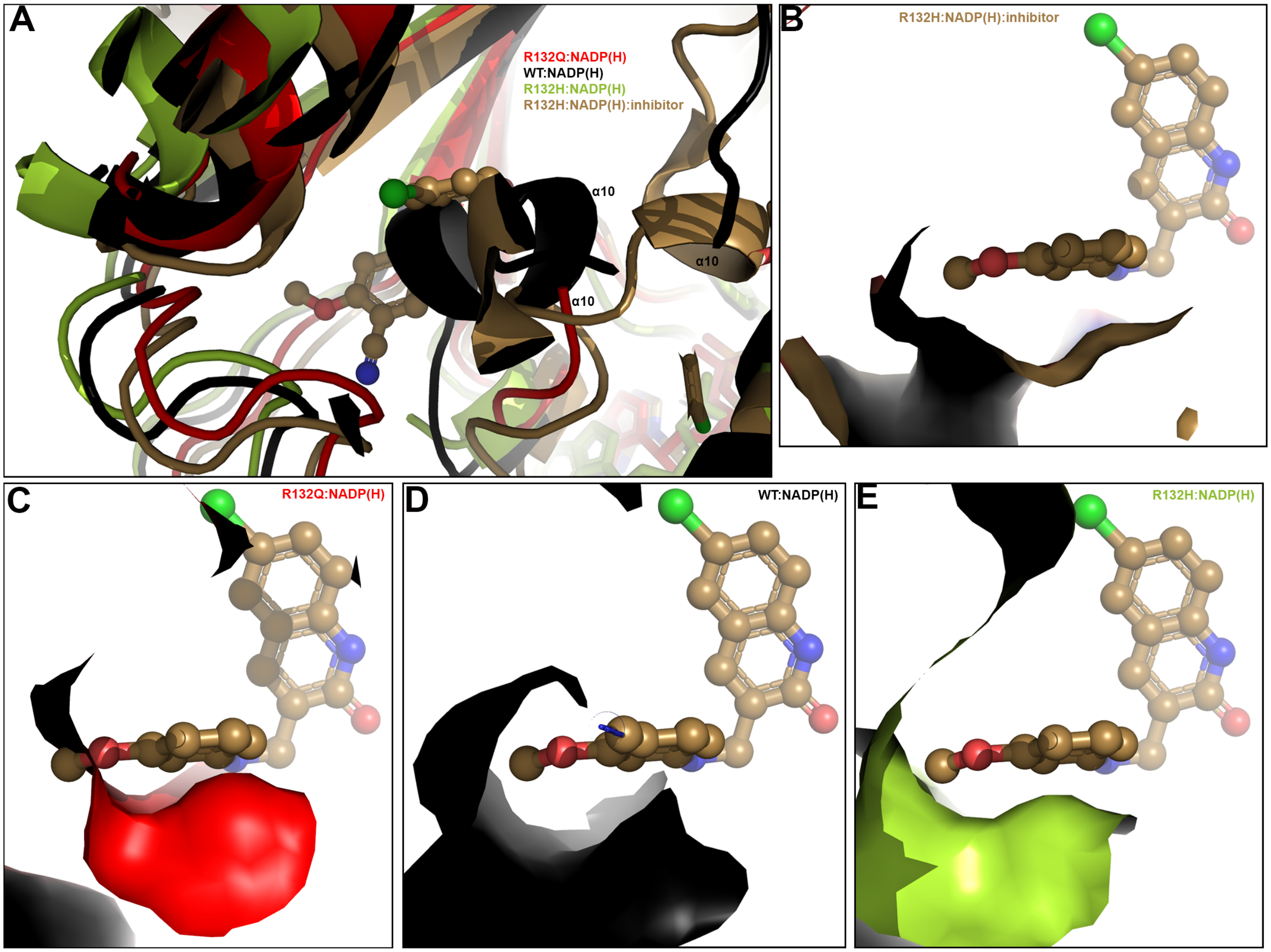


**Extended Data Fig. 10. Possible mechanisms of IDH1 R132Q resistance.** We have reported previously that IDH1 R132Q binds selective mutant IDH1 inhibitors poorly. A) A previously solved structure of a selective IDH1 R132H inhibitor (6O2Y) [4] was aligned to WT [1], R132H [3], and R132Q binary complexes. In (B-E), residues A111-V121 are shown as a surface. B) The structure of the inhibitor bound to a R132H:NADP(H) complex [4]. Residues A111-V121 in R132Q:NADP(H) (C) and in WT:NADP(H) (D) obstruct the inhibitor binding pocket. E) The inhibitor could be accommodated in the structure of R132H:NADP(H) [3].

### References


1. Xu, X. *et al.* Structures of human cytosolic NADP-dependent isocitrate dehydrogenase reveal a novel self-regulatory mechanism of activity. *J Biol Chem* **279**, 33946–57 (2004).
2. Rendina, A. R. *et al.* Mutant IDH1 enhances the production of 2-hydroxyglutarate due to its kinetic mechanism. *Biochemistry* **52**, 4563–77 (2013).
3. Yang, B., Zhong, C., Peng, Y., Lai, Z. & Ding, J. Molecular mechanisms of ‘off-on switch’ of activities of human IDH1 by tumor-associated mutation R132H. *Cell Res* **20**, 1188–200 (2010).
4. Lin, J. *et al.* Discovery and optimization of quinolinone derivatives as potent, selective, and orally bioavailable mutant isocitrate dehydrogenase 1 (mIDH1) inhibitors. *J Med Chem* **62**, 6575–6596 (2019).
