## Supplemental Material for "Active site remodeling in tumor-relevant IDH1 mutants drives distinct kinetic features and potential resistance mechanisms"

#### Supplementary Information Table of Contents:

Supplementary Figs. 1-6

Supplementary Tables 1-6

Supplementary References

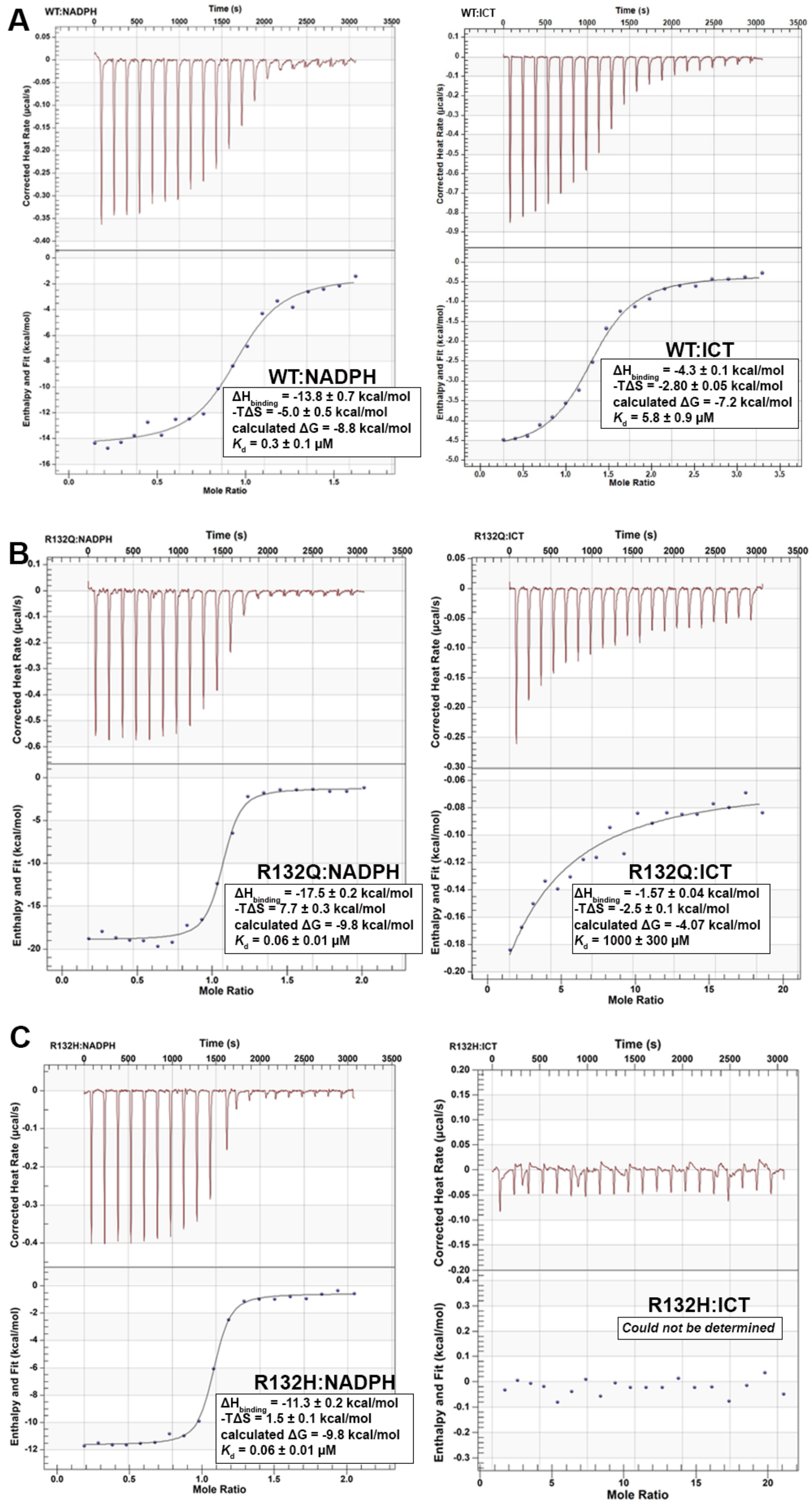


**Supplementary Fig. 1. Binding affinities of IDH1 WT, R132Q and R132H for NADPH and ICT using isothermal titration calorimetry.** A) IDH1 WT. B) IDH1 R132Q. C) IDH1 R132H.

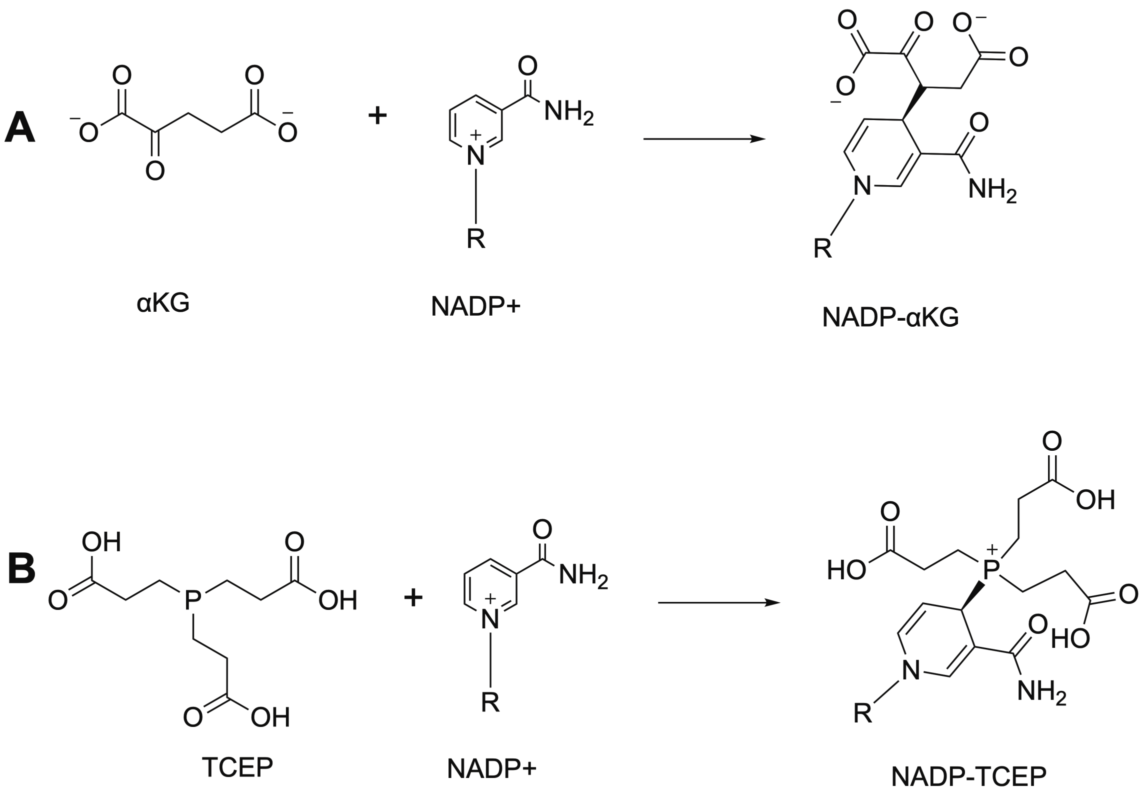


**Supplementary Fig. 2. Observed adduct formation upon IDH1 R132Q crystallization.** A) NADP-αKG adduct formation. B) NADP-TCEP adduct formation.

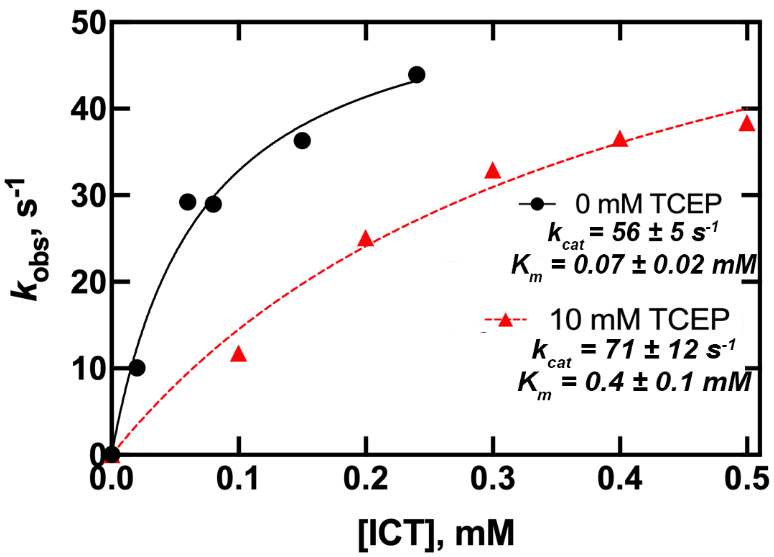


**Supplementary Fig. 3. TCEP treatment has less of an effect on IDH1 WT catalysis.** While conversion of ICT to αKG by IDH1 R132Q is associated with a 19-fold increase in $K_m$ in the presence of 10 mM TCEP, we see only a 6-fold increase for IDH1 WT.

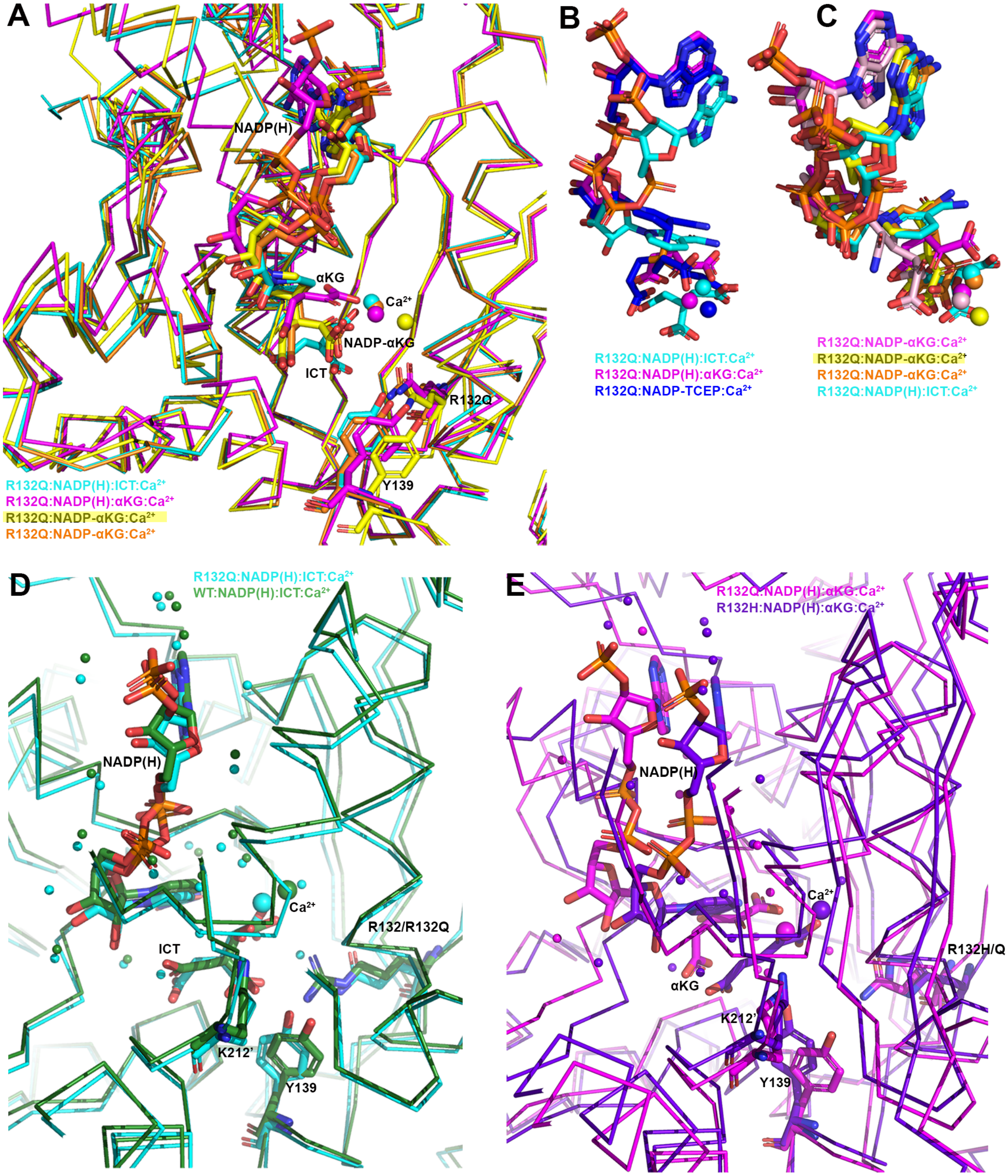


**Supplementary Fig. 4. IDH1 R132Q structures vary in substrate binding location, degree of nicotinamide ring puckering, and active site waters.** A) Monomer-based alignment of the R132Q:NADP-αKG:$Ca^{2+}$ monomer (orange) that aligns with R132Q:NADP(H):ICT:$Ca^{2+}$, and the R132Q:NADP-αKG:$Ca^{2+}$ monomer (yellow) that aligns as a transition between R132Q:NADP(H):ICT:$Ca^{2+}$ and R132Q:NADP(H):αKG:$Ca^{2+}$. B, C) Only the NADP(H) and ICT or αKG, or the NADP-adduct, with $Ca^{2+}$, are highlighted. Though the nicotinamide ring in the R132Q:NADPH:αKG:$Ca^{2+}$ structure could not be confidently modeled, comparisons of the planar nicotinamide ring of R132Q:NADP(H):ICT:$Ca^{2+}$ versus the puckered rings of the NADP-adducts are featured. Monomer-based alignment of R132Q:NADP(H):ICT:$Ca^{2+}$, R132Q:NADPH:αKG:$Ca^{2+}$, and R132Q:NADP-TCEP:$Ca^{2+}$ are shown in B). Monomer-based alignment of R132Q:NADP(H):ICT:$Ca^{2+}$, R132Q:NADPH:αKG:$Ca^{2+}$, and three observed R132Q:NADP-αKG:$Ca^{2+}$ monomers are shown in C). Dimer-based alignments are shown in (D) and (E). D) Water molecules (smaller spheres) within 3.5 Å of the substrates are highlighted in ICT-containing structures. E) Water molecules (smaller spheres) within 3.5 Å of the substrates are highlighted in αKG-containing structures.

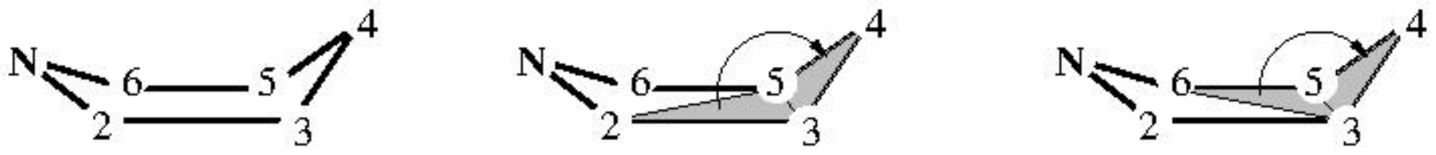


**Supplementary Fig. 5. Atom labeling and choice of dihedral angles for determination of deviation from planarity of atom C4.** The positions of the N atom 1 and the opposite C atom 4 are referenced to the plane defined by the roughly coplanar atoms 2, 3, 5, and 6. The average of the dihedral angles 2-3-5-4 and 6-3-5-4 shown here is subtracted from 180º to yield $\Delta\theta_{\mathrm{C}}$ as a metric for the deviation from planarity of C4, while the average of 3-2-6-1 and 5-2-6-1 subtracted from 180º is used to calculate $\Delta\theta_{\mathrm{N}}$ for N1.

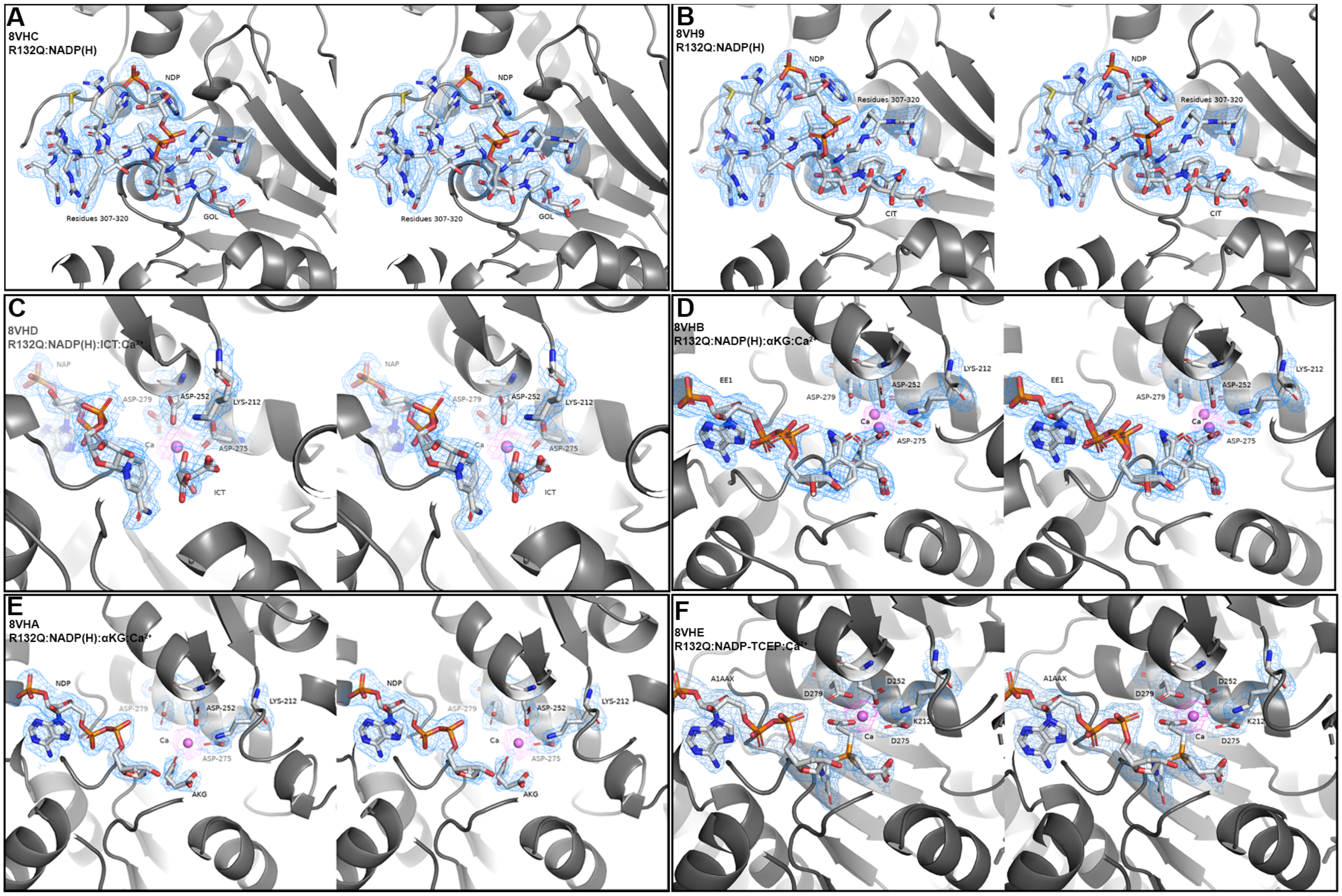


**Supplementary Fig. 6. Stereo-image of a section of the electron density map for the IDH1 R132Q structures.** A) R132Q:NADP(H) (PDB ID 8VHC), with residues 307-320, NADP(H) (NDP), and glycerol (GOL) highlighted. B) R132Q:NADP(H) (PDB ID 8VH9), with residues 307-320, NADP(H) (NDP), and citrate (CIT) highlighted. C) R132Q:NADP(H):ICT:$Ca^{2+}$ (PDB ID 8VHD), with D279, D252, K212, D275, NADP(H) (NAP), $Ca^{2+}$, and ICT highlighted. D) R132Q:NADP(H):αKG:$Ca^{2+}$ (PDB ID 8VHB), with D279, D252, K212, D275, NADP- αKG adduct (EE1), and $Ca^{2+}$ highlighted. E) R132Q:NADP(H):αKG:$Ca^{2+}$ (PDB ID 8VHA). F) R132Q:NADP-TCEP:$Ca^{2+}$ (PDB ID 8VHE), with D279, D252, K212, D275, NADP- TCEP adduct (A1AAX), and $Ca^{2+}$ highlighted.

**Supplementary Table 1. Active site and back cleft measurements assessing open versus closed conformations.** By convention, active site clefts are measured as the distance between residues 76 in chain A and 250’ in chain B, and back clefts are measured as the distance between residues 199 to 342 in chain A. Active, closed conformations are associated with small active site clefts and large back clefts.

| PDB ID | IDH1 complex (monomer/monomer listed if applicable) | I76 to L250’ active site cleft distance, Å | I76’ to L250 active site cleft distance, Å | M199 to H342 back cleft distance, Å | M199’ to H342’ back cleft distance, Å |
|---|---|---|---|---|---|
| 8VHC | R132Q:NADP(H) [a] | 14.7 | 16.9 | 9.1 | 8.9 |
| 8VH9 | R132Q:NADP(H) [b] | 14.2 | 17.1 | 8.7 | 8.4 |
| 8VHD | R132Q:NADP(H):ICT:$Ca^{2+}$/ R132Q:NADP(H):$Ca^{2+}$ | 8.8 (ICT)[c] | 10.4 (no ICT) | 10.1 (ICT) | 10.3 (no ICT) |
| 8VHB | R132Q:NADP(H):αKG:$Ca^{2+}$/ R132Q:NADP-αKG:$Ca^{2+}$ | 10.6 (αKG) | 11.7 (adduct) | 9.8 (αKG) | 9.5 (adduct) |
| 8VHA | R132Q:NADP(H):αKG:$Ca^{2+}$/ R132Q:NADP-αKG:$Ca^{2+}$ | 11.3 (αKG) | 12.2 (adduct) | 10.2 (αKG) | 10.0 (adduct) |
| 8VHA | R132Q:NADP-αKG:$Ca^{2+}$/ R132Q:NADP(H):$Ca^{2+}$ | 8.7 (adduct) | 11.8 (no αKG) | 9.7 (adduct) | 10.6 (no αKG) |
| 8VHE | R132Q:NADP-TCEP: $Ca^{2+}$ | 11.8 | 11.3 | 9.6 | 10.7 |
| 1T09[1] | WT:NADP(H) | 17.6 | 14.8 | 8.3 | 9.0 |
| 1T0L[1] | WT:NADP(H):ICT:$Ca^{2+}$ | 8.8 | 8.6 | 11.1 | 11.1 |
| 4L03[2] | G97D:NADP(H):αKG:$Ca^{2+}$ | 8.6 | 8.7 | 10.9 | 11.1 |
| 3MAR[3] | R132H:NADP(H) | 17.8 | 15.1 | 8.3 | 8.4 |
| 3MAP[3] | R132H:NADP(H):ICT | 16.1 | 14.1 | 7.7 | 8.0 |
| 4KZO[2] | R132H:NADP(H):αKG:$Ca^{2+}$ | 8.4 | 8.6 | 10.9 | 11.1 |

[a] Condition crystallized in sulfate
[b] Condition crystallized in citrate
[c] Dimers containing monomers with different molecules in the active site are indicated in the second column, and measurements associated with each monomer are identified by listing the active site molecule present in parenthesis.

**Supplementary Table 2. Relative energies and free energies of model $NADP^+$ and TCEP for binding, from B3LYP/pc-1 calculations.** The final energies and free energies reported were calculated by adding the counterpoise corrections to the energies of the geometries optimized with solvation.

| **Species** | **Relative $E_0$[a] (kcal mol$^{-1}$)** | **Relative $E$ (298K)[b] (kcal mol$^{-1}$)** | **Relative $G$ (kcal mol$^{-1}$)** |
|---|---|---|---|
| NADP + TCEP | 0 | 0 | 0 |
| NADP-TCEP TS | 2.5 | -2.2 | 24.6 |
| NADP-TCEP adduct | -4.3 | -9.4 | 20.0 |

[a] Relative electronic energy, including solvation and counterpoise corrections but excluding zero-point and thermal contributions.
[b] Relative energy, including solvation, counterpoise, zero-point, and thermal contributions.

**Supplementary Table 3. Dihedral angle measurements**. Using the angles shown in Supplementary Fig. 5, dihedral angles of the nicotinamide ring for NADP(H) or NADP-adduct molecules bound to IDH1 R132Q are shown. A sign convention has been applied such that if $\Delta\theta_{\mathrm{C}}$ and $\Delta\theta_{\mathrm{N}}$ have the same sign, the two corners of the ring bend away each other in chair fashion, whereas opposite signs indicate a boat-like conformation.

| **Species** | **Structural details** | $\Delta\boldsymbol{\theta}_{\mathbf{C}}$ **(º)** | $\Delta\boldsymbol{\theta}_{\mathbf{N}}$ **(º)** | **Conformation** |
|---|---|---|---|---|
| NADP(H) | R132Q:NADP(H), Fig 5A (chain A) | 2.4 | -4.1 | Planar |
| NADP(H) | R132Q:NADP(H), Fig 5A (chain B) | 4.8 | -7.6 | Planar |
| NADP(H) | R132Q:NADP(H):ICT:$Ca^{2+}$, Fig 6A (ICT monomer) | 0.4 | 0.2 | Planar |
| NADP(H) | R132Q:NADP(H):ICT:$Ca^{2+}$, Fig 6A (non-ICT monomer) | 0.3 | 0.1 | Planar |
| NADP-αKG<br><br>*DFT-calculated values:* | R132Q:NADP(H):αKG:$Ca^{2+}$, Fig 7A (adduct monomer opposite αKG monomer) | 25.0<br><br>*29* | -25.6<br><br>*-14* | Boat-like |
| NADP-αKG | R132Q:NADP(H):αKG:$Ca^{2+}$, Fig 7B (adduct monomer opposite αKG monomer) | 26.1 | -3.0 | Boat-like |
| NADP-αKG | R132Q:NADP(H):αKG:$Ca^{2+}$, Fig 7C (adduct monomer opposite non-αKG monomer) | 15.4 | -14.1 | Boat-like |
| NADP-TCEP<br><br>*DFT-calculated values:* | R132Q:NADP-TCEP:$Ca^{2+}$, Fig 7G (chain A) | 29.2<br><br>*25* | -1.1<br><br>*-11* | Boat-like |
| NADP-TCEP | R132Q:NADP-TCEP:$Ca^{2+}$, Fig 7G (chain B) | 26.0 | 9.3 | Chair-like |

**Supplementary Table 4. Steady-state kinetic parameters for conversion of ICT to αKG by IDH1 R132Q upon challenge with reducing agents**. TCEP treatment resulted in inhibition of the conventional reaction catalyzed by IDH1 R132Q. We did not observe any change in activity upon 10 mM TCEP treatment for the neomorphic reaction catalyzed by IDH1 R132H ($k_{cat}$ = 1.0 ± 0.03, $K_m$ = 0.42 ± 0.05) and by IDH1 R132Q ($k_{cat}$ = 2.11 ± 0.07, $K_m$ = 0.21 ± 0.04), which use NADPH as a substrate rather than $NADP^+$.

| **[Reducing agent]** | **$k_{cat,\ ICT \rightarrow \alpha KG}$ ($s^{-1}$)** | **$K_{M,\ ICT}$ (mM)** | **$k_{cat}/K_{M,\ ICT \rightarrow \alpha KG}$ ($mM^{-1}s^{-1}$)** |
|---|---|---|---|
| **[TCEP]** | | | |
| 0 mM | 2.1 ± 0.1 | 1.7 ± 0.5 | 1.2 ± 0.4 |
| 0.1 mM | 0.84 ± 0.04 | 1.6 ± 0.3 | 0.53 ± 0.09 |
| 0.5 mM | 2.3 ± 0.3 | 4 ± 1 | 0.7 ± 0.3 |
| 1 mM | 2.7 ± 0.3 | 6 ± 2 | 0.4 ± 0.1 |
| 2 mM | 3.1 ± 0.9 | 14 ± 8 | 0.2 ± 0.2 |
| 5 mM | 8 ± 4 | 83 ± 45 | 0.09 ± 0.07 |
| 10 mM | 2.7 ± 0.5 | 32 ± 9 | 0.09 ± 0.03 |
| **[DTT]** | | | |
| 0 mM | 2.1 ± 0.1 | 1.7 ± 0.5 | 1.2 ± 0.4 |
| 0.1 mM | 0.69 ± 0.05 | 0.9 ± 0.2 | 0.80 ± 0.02 |
| 0.5 mM | 2.4 ± 0.2 | 1.8 ± 0.8 | 1.3 ± 0.6 |
| 1 mM | 0.90 ± 0.06 | 2.1 ± 0.5 | 0.42 ± 0.09 |
| 2 mM | 2.2 ± 0.2 | 4 ± 1 | 0.62 ± 0.2 |
| 5 mM | 1.03 ± 0.06 | 1.1 ± 0.2 | 1.0 ± 0.2 |
| 10 mM | 2.0 ± 0.1 | 2.2 ± 0.7 | 0.9 ± 0.3 |
| **[BME]** | | | |
| 0 mM | 2.1 ± 0.1 | 1.7 ± 0.5 | 1.2 ± 0.4 |
| 0.1 mM | 4.2 ± 0.2 | 1.6 ± 0.3 | 2.5 ± 0.5 |
| 0.5 mM | 1.75 ± 0.07 | 1.3 ± 0.2 | 1.3 ± 0.2 |
| 1 mM | 2.0 ± 0.7 | 6 ± 7 | 0.34 ± 0.4 |
| 2 mM | 1.87 ± 0.08 | 2.5 ± 0.5 | 0.76 ± 0.2 |
| 5 mM | 4.9 ± 0.3 | 2.8 ± 0.5 | 1.7 ± 0.3 |
| 10 mM | 3.4 ± 0.3 | 5.9 ± 1.4 | 0.6 ± 0.2 |

#### Supplementary Table 5. HDX-MS parameters.

| Data Set | WT:NADP(H) | WT:NADP(H):ICT | WT:NADP(H):ICT:$Ca^{2+}$ |
|---|---|---|---|
| HDX reaction details | 50 mM Tris, 100 mM NaCl, pD= 7.85 @ 4 °C | 50 mM Tris, 100 mM NaCl, pD= 7.85 @ 4 °C | 50 mM Tris, 100 mM NaCl, pD= 7.85 @ 4 °C |
| HDX time course (min) | 0.5, 1, 2, 5 | 0.5, 1, 2, 5 | 0.5, 1, 2, 5 |
| HDX control samples | Disordered section of WT NT protein | Disordered section of WT NT protein | Disordered section of WT NT protein |
| Back-exchange (mean/IQR) | 42%/5% | 42%/5% | 42%/5% |
| # of Peptides | 112 | 112 | 112 |
| Sequence coverage | 99.8% | 99.8% | 99.8% |
| Average peptide length/Redundancy | 15.9/ 4.12 | 15.9/ 4.12 | 15.9/ 4.12 |
| Replicates (biological or technical) | 3 (technical) | 3 (technical) | 3 (technical) |
| Repeatability | 0.046 (average standard deviation) | 0.047 (average standard deviation) | 0.053 (average standard deviation) |
| Significant differences in HDX (Δ HDX > X D) | 0.25D (99% CI) | 0.25D (99% CI) | 0.25D (99% CI) |

| Data Set | R132H:NADP(H) | R132H:NADP(H):αKG | R132H:NADP(H):αKG:$Ca^{2+}$ |
|---|---|---|---|
| HDX reaction details | 50 mM Tris, 100 mM NaCl, pD= 7.85 @ 4 °C | 50 mM Tris, 100 mM NaCl, pD= 7.8 5@ 4 °C | 50 mM Tris, 100 mM NaCl, pD= 7.85 @ 4 °C |
| HDX time course (min) | 0.5, 1, 2, 5 | 0.5, 1, 2, 5 | 0.5, 1, 2, 5 |
| HDX control samples | Disordered section of WT NT protein | Disordered section of WT NT protein | Disordered section of WT NT protein |
| Back-exchange (mean / IQR) | 31%/5% | 31%/5% | 31%/5% |
| # of Peptides | 112 | 112 | 112 |
| Sequence coverage | 99.8% | 99.8% | 99.8% |
| Average peptide length / Redundancy | 15.9/ 4.12 | 15.9/ 4.12 | 15.9/ 4.12 |
| Replicates (biological or technical) | 3 (technical) | 3 (technical) | 3 (technical) |
| Repeatability | 0.089 (average standard deviation) | 0.105 (average standard deviation) | 0.079 (average standard deviation) |
| Significant differences in HDX ( Δ HDX > X D) | 0.25D (99% CI) | 0.25D (99% CI) | 0.25D (99% CI) |

| Data Set | R132Q:NADP(H) | R132Q:NADP(H):ICT | R132Q:NADP(H):ICT:$Ca^{2+}$ | R132Q:NADP(H):αKG | R132Q:NADP(H):αKG:$Ca^{2+}$ |
|---|---|---|---|---|---|
| HDX reaction details | 50 mM Tris, 100 mM NaCl, pD= 7.85 @ 4 °C | 50 mM Tris, 100 mM NaCl, pD= 7.85 @ 4 °C | 50 mM Tris, 100 mM NaCl, pD= 7.85 @ 4 °C | 50 mM Tris, 100 mM NaCl, pD= 7.85 @ 4 °C | 50 mM Tris, 100 mM NaCl, pD= 7.85 @ 4 °C |
| HDX time course (min) | 0.5, 1, 2, 5 | 0.5, 1, 2, 5 | 0.5, 1, 2, 5 | 0.5, 1, 2, 5 | 0.5, 1, 2, 5 |
| HDX control samples | Disordered section of WT NT protein | Disordered section of WT NT protein | Disordered section of WT NT protein | Disordered section of WT NT protein | Disordered section of WT NT protein |
| Back-exchange (mean / IQR) | 51% / 5% | 51% / 5% | 51% / 5% | 51% / 5% | 51% / 5% |
| # of Peptides | 112 | 112 | 112 | 112 | 112 |
| Sequence coverage | 99.8% | 99.8% | 99.8% | 99.8% | 99.8% |
| Average peptide length / Redundancy | 15.9/4.12 | 15.9/4.12 | 15.9/4.12 | 15.9/4.12 | 15.9/4.12 |
| Replicates (biological or technical) | 3 (technical) | 3 (technical) | 3 (technical) | 3 (technical) | 3 (technical) |
| Repeatability | 0.060 (average standard deviation) | 0.064 (average standard deviation) | 0.061 (average standard deviation) | 0.144 (average standard deviation) | 0.069 (average standard deviation) |
| Significant differences in HDX ( Δ HDX > X D) | 0.25D (99% CI) | 0.25D (99% CI) | 0.25D (99% CI) | 0.25D (99% CI) | 0.25D (99% CI) |

#### Supplementary Table 6. Crystallography parameters.

| ***Data collection*** | **R132Q:NADP(H) (sulfate condition)** | **R132Q:NADP(H) (citrate condition)** | **R132Q:NADP(H):ICT:$Ca^{2+}$** | **R132Q:NADP(H):αKG: $Ca^{2+}$** | **R132Q:NADP(H):αKG: $Ca^{2+}$** | **R132Q:NADP-TCEP:$Ca^{2+}$** |
|---|---|---|---|---|---|---|
| PDB code | 8VHC | 8VH9 | 8VHD | 8VHB | 8VHA | 8VHE |
| Space Group | P 43 21 2 | P 43 21 2 | P 1 21 1 | P 1 21 1 | P 1 21 1 | P 1 21 1 |
| Cell Dimensions<br>a, b, c (Å)<br>α, β, γ (°) | 82.884 82.884 303.926<br>90 90 90 | 81.085 81.085 306.136<br>90 90 90 | 84.408 103.894 108.272<br>90 98.539 90 | 84.404 105.807 109.782<br>90.00 98.443 90.00 | 83.821 104.86 107.711<br>90.00 98.188 90.00 | 84.298 107.348 109.941<br>90 99.187 90 |
| Resolution (Å) | 2.44 | 2.13 | 2.38 | 1.89 | 2.28 | 2.16 |
| Observations | 80846 (7762) | 518814 (53176) | 142446 (13835) | 288375 (28444) | 162883 (15717) | 304736 |
| Source | APS 24-ID-E | APS 24-ID-E | SSRL Beamline 12-1 | SSRL Beamline 12-1 | SSRL Beamline 12-1 | APS 24-ID-E |
| Wavelength (Å) | .979180 | .979180 | .97946 | .97946 | .97946 | .979180 |
| <I/σ(I)> | 22.37 | 18.29 | 5.28 | 12.71 | 10.65 | 13.0 |
| Completeness | 99.60 | 99.91 | 96.88 | 97.12 | 98.28 | 98.01 |
| ***Refinement*** | | | | | | |
| Resolution range (Å) | 72.76 - 2.44 (2.527 - 2.44) | 76.53 - 2.13 (2.206 - 2.13) | 39.18 - 2.38 (2.465 - 2.38) | 39.63 - 1.89 (1.958 - 1.89) | 39.15 - 2.28 (2.361 - 2.28) | 83.22 - 2.16 (2.237 - 2.16) |
| Unique reflections | 40453 (3891) | 58370 (5722) | 72946 (6517) | 148111 (14777) | 82712 (8073) | 101523 (10184) |
| Protein residues | 794 | 795 | 1650 | 1653 | 1649 | 1660 |
| Ligand atoms | 186 | 199 | 237 | 448 | 334 | 327 |
| Water Atoms | 75 | 343 | 801 | 1069 | 462 | 933 |
| R-work | 0.1922 (0.3187) | 0.1756 (0.2863) | 0.1694 (0.2144) | 0.1680 (0.2685) | 0.1673 (0.2185) | 0.1732 (0.2291) |
| R-free | 0.2398 (0.3678) | 0.2153 (0.3172) | 0.2199 (0.2885) | 0.1723 (0.2692) | 0.2222 (0.2925) | 0.2218 (0.2770) |
| Wilson B-Factor | 58.0 | 41.1 | 22.9 | 29.1 | 35.6 | 34.1 |
| RMS (bonds) | 0.004 | 0.010 | 0.002 | 0.008 | 0.009 | 0.005 |
| RMS (angles) | 0.59 | 1.02 | 0.42 | 0.92 | 0.91 | 0.67 |
| Clash Score | 12.42 | 7.39 | 3.29 | 9.69 | 8.02 | 3.36 |
| ***Ramachandran plot*** | | | | | | |
| Favored (%) | 95.78 | 95.68 | 96.59 | 96.84 | 96.59 | 96.49 |
| Allowed (%) | 4.09 | 4.32 | 3.29 | 3.04 | 3.35 | 3.51 |

### References


1. Xu, X. *et al.* Structures of human cytosolic NADP-dependent isocitrate dehydrogenase reveal a novel self-regulatory mechanism of activity. *J Biol Chem* **279**, 33946–57 (2004).
2. Rendina, A. R. *et al.* Mutant IDH1 enhances the production of 2-hydroxyglutarate due to its kinetic mechanism. *Biochemistry* **52**, 4563–77 (2013).
3. Yang, B., Zhong, C., Peng, Y., Lai, Z. & Ding, J. Molecular mechanisms of ‘off-on switch’ of activities of human IDH1 by tumor-associated mutation R132H. *Cell Res* **20**, 1188–200 (2010).
